# Chronic Wound Milieu Challenges Essential Oils’ Antibiofilm Activity

**DOI:** 10.1101/2023.06.21.545846

**Authors:** Malwina Brożyna, Weronika Kozłowska, Katarzyna Malec, Justyna Paleczny, Jerzy Detyna, Krystyna Fabianowska-Majewska, Adam Junka

## Abstract

The treatment of infected non-healing wounds poses a significant challenge to contemporary medicine. Essential oils (EOs) are being increasingly investigated as potential antibiofilm agents for the management of biofilm-related wound infections. However, their *in vitro* antimicrobial activity reported in literature does not necessarily reflect the actual *in vivo* activity due to, among others, methodological imperfections of performed tests. In our study, we cultivated a *Staphylococcus aureus* biofilm in a novel IVWM (In Vitro Wound Milieu) medium designed to simulate the wound environment and compared it with biofilms cultured in the standard microbiological Tryptic Soy Broth (TSB) medium. We examined and compared critical biofilm properties such as morphology, biomass, metabolic activity, cell count, thickness, and spatial distribution of live and dead cells. Subsequently, staphylococcal biofilms and planktonic cells cultured in both media were exposed to the activity of thyme or rosemary EOs (T-EO, R-EO, respectively). We found that morphology of biofilms cultured in IVWM resembled more the morphology of biofilms visualized in the non-healing wounds than biofilms cultured in TSB. The biomass, metabolic activity, cell number, and the ratio of live to dead cells of *S. aureus* biofilms were all lower in IVWM compared to the TSB medium. Additionally, while EOs demonstrated overall significant anti-staphylococcal activity, their efficacy varied depending on the medium used. Generally, EOs displayed lower antimicrobial activity (against planktonic cells) in the IVWM than in the TSB medium. Interestingly, T-EO caused a higher reduction of biofilm cells in IVWM than in the TSB medium, in contrast to R-EO. Our findings suggest that EOs hold promise as agents for the treatment of biofilm-related wound infections. However, it is crucial to apply *in vitro* conditions that closely reflect the wound infection site to gain an accurate insight into the real-world activity of these antimicrobial/antibiofilm agents.

## 1 Introduction

The dissemination of antibiotic-resistant microorganisms poses a global threat to public health care. As the efficacy of existing antibiotics decreases and new ones remain undeveloped, the applicability of nonantibiotic therapeutics to combat infections is being thoroughly investigated worldwide (Mühlen and Dersch, 2016).

Non-healing wounds are particularly susceptible to infections caused by antibiotic-resistant microorganisms (Bowler, 2018). These wounds affect 20 million patients annually and require more than 31 billion USD per year for treatment (Leaper et al., 2015). The five-year mortality rate for people with diabetic non-healing wounds is comparable to the five-year mortality rate for patients with cancer (30.5% vs. 31%, respectively) (Armstrong et al., 2020).

Infections, one of the most frequent complications of non-healing wounds, are caused by biofilms – complex microbial communities embedded within an extracellular matrix (ECM) (Percival et al., 2012). This scaffold provides not only a physical framework for the biofilm but also impedes the immune response and penetration of antimicrobial agents. Furthermore, some microbial cells within the biofilm matrix exhibit low cellular activity, rendering them insensitive to antibiotics that target cellular transcription or translation processes (Povolotsky et al., 2021). In general, biofilms exhibit significantly higher tolerance to the immune system and antimicrobials compared to their free-floating (planktonic) counterparts.

Thus, the treatment of infected non-healing wounds is a major challenge for modern medicine. Current strategies to eradicate wound biofilms involve the debridement (removal) of infected tissue, combined with the topical application of dressings and antiseptic agents (Kaiser et al., 2021). Topical administration of antibiotics is no longer recommended due to their low activity against biofilms and the potential to induce microbial resistance, hypersensitivity, or contact allergy (Siddiqui and Bernstein, 2010).

Modern antiseptic agents are considered first-line antimicrobials for non-healing wound management due to their broad spectrum of antimicrobial activity, non-specific mode of action, and low *in vivo* cytotoxicity (Alves et al., 2021). However, several reports indicate the potential for microorganisms to develop tolerance to modern antiseptic agents (compounds obtained through industrial chemical synthesis). The survival of *Burkholderia cepacia* in octenidine dihydrochloride-containing solutions and the increased tolerance of certain strains of *Pseudomonas aeruginosa* to octenidine dihydrochloride exemplify such risks (Becker et al., 2018; Shepherd et al., 2018)

It can be inferred that the effectiveness of systemic antibiotic therapy against wound biofilms is limited, and the topical administration of these antimicrobials is not recommended. The use of modern antiseptic agents (e.g., polyhexanide, povidone-iodine) still correlates with favorable clinical outcomes, but the example of octenidine dihydrochloride raises concerns that the efficacy of these antiseptics may also be reduced due to microorganisms’ tendency to develop resistance (Becker et al., 2018; Shepherd et al., 2018). Therefore, not only non-antibiotic but also non-antiseptic approaches are now considered next-generation strategies for treating biofilm-related infections. These methods include the use of natural compounds, enzymes, and other bioactive molecules that can disrupt biofilm formation and/or enhance host immune responses (Silva et al., 2020). In this regard, plant-derived essential oils (EOs) are of great interest as potential antibiofilm agents. These multi-component volatile liquids exhibit a broad spectrum of antimicrobial activity against Gram-positive and Gram-negative bacteria and fungi (Agreles et al., 2021). EOs can also interfere with or impede various processes occurring in biofilms, such as adhesion, quorum-sensing, or modulation of the expression of biofilm-related genes (Reichling, 2020).

The non-specific mode of EOs’ antimicrobial action is considered to limit the risk of triggering bacterial resistance (Yap et al., 2014). Thanks to their low cytotoxicity, anti-inflammatory activity, and ability to promote the wound-healing process, the use of EOs can be perceived as an effective future strategy for the treatment of non-healing wound infections (Costa et al., 2019). Although several *in vitro* studies report significant antimicrobial properties of EOs, these likely do not reflect their actual *in vivo* activity (Orchard et al., 2017; Sienkiewicz et al., 2017; Patterson et al., 2019). Major shortcomings of this *in vitro,* research include using standard microbiological media (rather than media reflecting the composition of wound exudate) and the evaluation of EOs’ activity primarily against planktonic forms of microbes, not against biofilms. Recent reports indicate that providing a milieu that mimics the wound environment in *in vitro* tests significantly alters key biofilm characteristics such as metabolic activity, three-dimensional structure, and matrix composition, thereby affecting its tolerance to antimicrobial agents (Thaarup and Bjarnsholt, 2021; Vyas et al., 2022). Therefore, in this study, we cultivated a *Staphylococcus aureus* biofilm (one of the main factors of wound infection) in a newly formulated medium known as the IVWM (In Vitro Wound Milieu), which comprises serum, cell-matrix elements, and host factors that reflect the wound environment (Kadam et al., 2021). After conducting a thorough analysis of the key properties of the biofilm grown in IVWM and comparing them to those of the biofilm cultivated in standard microbiological TSB medium, we exposed staphylococcal biofilms to the activity of thyme and rosemary EOs (T-EO, R-EO, respectively). We also carried out control analyses on planktonic *Staphylococcus aureus* cells. To the best of our knowledge, this is the first study that assesses the antimicrobial properties of EOs under conditions that resemble a chronic wound milieu.

## 2 Materials and methods

### 2.1 Microorganisms

One reference strain, *Staphylococcus aureus* ATCC 6538 (American Type and Culture Collection), and eleven clinical isolates were tested for research purposes. The clinical strains included four MSSA strains (Methicillin-Susceptible *Staphylococcus aureus*) and seven MRSA strains (Methicillin-Resistant *Staphylococcus aureus*). The MSSA strains were marked S3, S6, S8, S11; MRSA as R1, R8-R13. The strains are part of the Strain and Line Collection of the Pharmaceutical Microbiology and Parasitology Department of the Medical University of Wroclaw.

### 2.2 Essential oils

The antimicrobial activity of two commercial essential oils (EOs) was tested:

- thyme oil, thyme chemotype (T-EO, obtained from *Thymus vulgaris* L. leaves), produced by Instytut Aromaterapii, Poland;
- rosemary oil, camphor chemotype (R-EO, obtained from *Rosmarinus officinalis* L. leaves), produced by Instytut Aromaterapii, Poland.

### 2.3 Culture conditions

Bacteria were cultured in two media:

1. Standard microbiological Tryptic Soy Broth (Biomaxima, Poland) marked TSB. The detailed composition of TSB is presented in **Table 1**.
2. Medium prepared according to the formula presented by (Kadam et al., 2021), marked IVWM (In Vitro Wound Milieu).

**Table 1.** Composition of Tryptic Soy Broth (TSB) according to the manufacturer’s specification **(A)**. Composition of In Vitro Wound Milieu (IVWM) **(B)**.

| (A) TSB |  |
| --- | --- |
| Pancreatic digest of casein | 17 g/L |
| Peptic digest of soybean | 3 g/L |
| Dipotassium hydrogen phosphate | 2.5 g/L |
| Sodium chloride | 5 g/L |
| Glucose monohydrate | 2.5 g/L |
| (B) IVWM |  |
| Fetal Bovine Serum | 70% |
| Fibronectin | 30-60 µg/mL |
| Fibrinogen | 200-400 µg/mL |
| Lactoferrin | 20-30 µg/mL |
| Lactic acid | 11-12 mM |
| Collagen | 10-12 µg/mL |
| Saline | 19.6% |

Sterile Fetal Bovine Serum (Biowest, France, cat. no. S181H) was used as the base component of IVWM. Firstly, stock solutions of the components were prepared as follows:

- fibronectin (Human plasma fibronectin, Sigma-Aldrich, USA, cat. no. FC010) 1 mg/mL solution in autoclaved distilled water,

- fibrinogen (Fibrinogen from human plasma, Sigma-Aldrich, USA, cat. no. F3879) 10 mg/mL solution in saline (Stanlab, Poland),

- lactoferrin (Lactoferrin human, Sigma-Aldrich, USA, cat. no. L4040) 2 mg/mL solution in Dulbecco’s Phosphate Buffered Saline (Sigma-Aldrich, USA),

- lactic acid (Sigma-Aldrich, USA, cat. no. W261114) 11.4 M solution,

and they were filtered using a 0.22 μm syringe filter (Sungo, Europe). Collagen (Collagen solution from bovine skin, concentration 2.9-3.2 mg/mL, Sigma-Aldrich, USA, cat. no. C4243) was purchased sterile. The IVWM was obtained by combining the ingredients at concentrations presented in **Table 1**. The medium was stored for a maximum of seven days at 2-8°C and was protected from light.

### 2.4 Assessment of EOs Chemical Composition using Gas Chromatography-Mass Spectrometry (GC-MS)

EOs were diluted 50 times with hexane (JTB, GB), vortexed, and immediately analyzed. The analysis was performed on Agilent 7890B GC system coupled with 7000GC/TQ connected with PAL RSI85 autosampler (Agilent Technologies, USA) and equipped with an HP-5 MS column (30 m × 0.25 mm × 0.25 μm). Helium was used as a carrier gas at a total flow of 1 mL/min, and the injection mode was split in a ratio of 1:100. Analysis conditions were as follows: the initial temperature was 50°C for 1 min, then increased to 4°C/min to 170°C, and then 10°C/min to 280°C which maintained for 2 min. The MS detector settings were as follows: ionization voltage 70 eV, transfer line, source, and quadrupole temperature – 320, 230, and 150°C, respectively. Detection was performed in full scan mode in a range of 30-400 m/z. Identification was based on a comparison of retention index (RI) and mass spectra with data from the NIST 17.1 library and literature. Linear retention indexes were determined using a mixture of C8-C20 saturated alkanes (Sigma-Aldrich, USA) under the same conditions as for EOs. The relative abundance of each EO component was expressed as percentage content based on peak area normalization. The MassHunter Workstation Software version B.09.00 was used for peak normalization. All analyses were performed in triplicate.

### 2.5 Evaluation of biofilm biomass, biofilm metabolic activity, and the number of colony-forming units

Biofilm mass and biofilm metabolic activity of one reference and eleven clinical *S. aureus* strains were evaluated in TSB (Tryptic Soy Broth, Biomaxima, Poland) or IVWM (In Vitro Wound Milieu) medium. The number of colony-forming units was assessed for the reference strain. For this purpose, the bacteria were pre-incubated overnight in an appropriate medium (TSB or IVWM) at 37°C. Next, the bacterial suspensions were prepared in saline (Stanlab, Poland) and adjusted to 0.5 MF (McFarland, 1.5 × 10^^8^ CFU/mL (Colony-Forming Unit) using a densitometer (DEN-1, Biosan SIA, Latvia), and diluted thousand times in TSB or IVWM. 500 µL of this suspension was poured into the wells of a 48-well polystyrene plate (Wuxi Nest Biotechnology, China) and incubated for 24 h under static conditions at 37°C. To assess the total biofilm mass, crystal violet staining was performed. The level of biofilm activity was evaluated using tetrazolium staining. Both tests were performed in two independent experiments in six replicates. Quantitative culturing was performed in one experiment in triplicate to determine the number of colony-forming units.

- Evaluation of biofilm biomass level using crystal violet assay

After the biofilm culturing described above, the medium was removed, and the plates were dried for 10 min (37°C). Subsequently, 500 µL of 20% (v/v) crystal violet (Aqua-med, Poland) solution in water was added to the wells, and the plates were kept at room temperature for 10 min. The stain was gently removed, the biofilm was washed once with 500 µL of saline (Chempur, Poland), and the plates were incubated at 37°C for 10 min. Next, violet crystals were dissolved with 500 µL of 30% (v/v) acetic acid (Stanlab, Poland) water solution, and the plates were shaken for 30 min at 450 rpm (Mini-shaker PSU-2T, Biosan SIA, Latvia) at room temperature. 100 µL of the solution was transferred from one well in four replicates to 96-well plates (Wuxi Nest Biotechnology, China), and the absorbance was measured at 550 nm using a spectrophotometer (Multiskan Go, Thermo Fisher Scientific, USA). The average absorbance was calculated for each sample. The absorbance of media without bacteria was also measured, and their average values were subtracted from the absorbance of each sample. Based on the results, the strains were divided into four groups according to their biofilm biomass:

- high biofilm biomass in TSB: S11, R12, R13;
- low biofilm biomass in TSB: ATCC 6538, S3, R1;
- high biofilm biomass in IVWM: S6, S8, R9;
- low biofilm biomass in IVWM: R8, R10, R11.
- Assessment of biofilm activity level using tetrazolium staining

The biofilm was cultured as presented above, and the medium was removed from above the cells. Next, metabolically active biofilm cells were two-hours stained with 500 µL of 0.1% (w/v) TTC solution (2,3,5-triphenyl-tetrazolium chloride, Sigma-Aldrich, USA) in TSB (Tryptic Soy Broth, Biomaxima, Poland) or IVWM (In Vitro Wound Milieu) medium at 37°C. The medium was pipetted-out, and the plates were dried for 10 min at 37°C. 500 µL of methanol (Stanlab, Poland) and acetic acid (Stanlab, Poland) (9:1 ratio) solution was introduced to the wells, and the plates were shaken (Mini-shaker PSU-2T, Biosan SIA, Latvia) for 30 min at room temperature (400 rpm). 100 µL of the solution was transferred from one well in four replicates to 96-well plates (Wuxi Nest Biotechnology, China), and the absorbance was measured at 490 nm using a spectrophotometer (Multiskan Go, Thermo Fisher Scientific, USA). The average absorbance was calculated for each sample. The absorbance of media without bacteria was also measured, and their average values were subtracted from the absorbance of each sample.

- Assessment of the number of colony-forming units

As described above, the biofilm was cultured in polystyrene plates in TSB or IVWM medium. Subsequently, the medium was removed, and each well was shaken with 500 µL of 0.1% (w/v) saponin (VWR Chemicals, USA) water solution for 30 s at 600 rmp. Solutions from each well were resuspended and transferred to Eppendorf tubes. Plates were shaken again for 30 s/600 rpm with the fresh saponin solution (500 µL), and solutions from both steps were combined. The serial dilutions of the suspension were then prepared in saline solution and cultured onto Mueller–Hinton agar (Biomaxima, Poland) Petri dishes (Noex, Poland) and incubated for 24 h at 37°C. The CFU number was counted using ImageJ (National Institutes of Health, Bethesda, MD, USA, accessed on 1 December 2022).

### 2.6 Visualization of live and dead biofilm-forming cells using fluorescent dyes and a Confocal Microscopy

The staphylococcal strains were cultured in TSB (Tryptic Soy Broth, Biomaxima, Poland) or IVWM (In Vitro Wound Milieu) medium in 24-well plates (Wuxi Nest Biotechnology, China). The preparation of suspensions and biofilm cultivation conditions were the same as described in the section “Evaluation of biofilm biomass, biofilm metabolic activity, and the number of colony-forming units”. Filmtracer™ LIVE/DEAD™ Biofilm Viability Kit (Thermo Fischer Scientific, USA), prepared according to the manufacturer’s instruction, was applied as a dye to assess the membrane integrity and indirectly to measure a relative number of live and dead biofilm-forming cells and also to visualize the morphology/spatial distribution of cells within biofilm structure. The microscopic visualizations were performed using an SP8 MP laser-scanning confocal microscope (Leica, Germany). SYTO-9 showing live bacteria was excited at 488 nm wavelength using a laser line (SP8). The collected emission was within the 500–530 nm range. Propidium iodide (PI) for the visualization of dead bacteria was a 552 nm laser line (SP8). The emission of PI was collected within the 575–625 nm SP8 ranges. The acquisition was performed using 20µm dry objectives in a sequence to avoid a spectral bleed through. For a given set of experimental conditions (untreated biofilm and biofilms with EOs), the same acquisition settings were applied to each system to enable quantitative comparisons between the conditions. The settings and signal intensity were always set on the brightest samples to avoid oversaturation. Next, the pictures were processed using ImageJ (National Institutes of Health, Bethesda, MD, USA) software. The whole captured picture was treated as the Region of Interest (ROI). The mean grey value (MGV) of each ROI was recorded for green and red fluorescence channels and served as an estimator of changes in the number of live and dead cells, respectively. The MGV was considered the sum of gray values of all the pixels in the selection divided by the number of pixels. For RGB images recorded for the purpose of this analysis, the MGV was calculated by converting each pixel to grayscale using the Equation: gray = 0.299red + 0.587green + 0.114blue.

### 2.7 Visualization of biofilm using a Scanning Electron Microscope

The *S.aureus* ATCC 6538 biofilm strain was visualized with a Scanning Electron Microscope (SEM, Auriga 60, ZEISS, Germany). Firstly, 1 mL of 2% (w/v) Bacteriological Lab Agar (Biomaxima, Poland) was poured into a 24-well plate (Wuxi Nest Biotechnology, China) and left for solidification. Next, 500 µL of the appropriate medium (TSB (Tryptic Soy Broth, Biomaxima, Poland) or IVWM (In Vitro Wound Milieu)) was poured into the wells with agar, and the plate was kept refrigerated for 24 h. Subsequently, the medium was removed from above the agar, and the bacterial suspension in TSB or IVWM medium was prepared as described in the section “Evaluation of biofilm biomass, biofilm metabolic activity, and the number of colony-forming units”, and added in the amount of 500 µL to the agar wells. The plate was incubated for 24 h at 37°C under static conditions. The medium was then removed, and 500 µL of 4.5% (v/v) glutaraldehyde (ChemPur, Poland) was poured. Next, samples were dried in a critical point dryer EM CPD300 (Leica Microsystems, Germany). Subsequently, the samples were subjected to sputtering with Au/Pd (60:40) using EM ACE600, Leica sputter (Leica Microsystems, Germany). The sputtered samples were examined using a scanning electron microscope (SEM, Auriga 60, Zeiss, Germany).

### 2.8 Evaluation of Minimal Inhibitory and Minimal Biofilm Eradication Concentrations of EOs Emulsions

The antimicrobial activity of T-EO or R-EO was assessed in 96-well plates (Wuxi Nest Biotechnology, China) in TSB (Tryptic Soy Broth, Biomaxima, Poland) or IVWM (In Vitro Wound Milieu) media. Tests were carried out in two independent experiments in three replicates. For both experiments, bacterial suspension was prepared as follows. From overnight cultures in TSB or IVWM medium suspensions were prepared in saline (Stanlab, Polnad) and adjusted to 0.5 MF (McFarland, 1.5 × 10^^8^ CFU/mL (Colony-Forming Unit)) using a densitometer (DEN-1, Biosan SIA, Latvia), and diluted a thousand times in TSB or IVWM. EOs were tested as emulsions in TSB or IVWM medium with the addition of Tween 20 (Pol-aura, Poland). In the T-EO emulsions stock solution, the oil constituted 5% (v/v) of the emulsion volume, and in R-EO, 10% (v/v). Tween 20 constituted 1% (v/v) of the entire volume on both emulsions. The emulsions were prepared as follows: the EOs were mixed with Tween 20 for 30 min using a magnetic stirrer (IKA RH basic 2, IKA-Werke GmbH & CO. KG, Deutschland). Next, the medium was gradually added and stirred for one hour more at room temperature.

- Minimal Inhibitory Concentration (MIC) evaluation

To evaluate MIC values, geometric concentrations of EOs’ stock solutions were prepared in TSB or IVWM medium at concentrations ranging from 5 % to 0.002% (v/v) for T-EO and from 10% to 0.005% (v/v) for R-EO and added in a volume of 100 µL to the wells of the plates. Next, 100 µL of bacterial suspensions in TSB or IVWM medium (prepared as described above) were added to the wells, and the plates were incubated for 24 hours at 37°C with shaking (Mini-shaker PSU-2T, Biosan SIA, Latvia) at 450 rpm. Controls of bacterial growth (bacteria in TSB or IVWM medium) and medium sterility (medium only) were also prepared. After incubation, 20 µL of 1% (w/v) TTC solution (2,3,5-triphenyl-tetrazolium chloride, Sigma-Aldrich, USA) in the medium was added to each well, and plates were incubated for 2 hours under the same conditions. MIC values were evaluated in the first well, where no red color was observed. The influence of five concentrations [(%) (v/v)] of Tween 20 on *S. aureus* ATCC 6538 planktonic forms cultured in TSB medium was also evaluated in the same manner as the test with EOs. In addition, the absorbance of the samples at 580 nm was measured using a spectrophotometer (Multiskan Go, Thermo Fisher Scientific, USA) before adding the TTC solution. The percentage of cell viability was calculated in each Tween 20 concentration with respect to the growth control. The test was performed in one experiment in six replicates.

- Minimal Biofilm Eradication Concentration (MBEC) evaluation

To analyze the antibiofilm properties of EOs emulsions, biofilms were first cultivated in polystyrene plates. For this purpose, 100 µL of TSB or IVWM was added to the wells of the plates, and 100 µL of bacterial suspensions in TSB or IVWM medium (prepared as described above) was poured. The plates were incubated at 37°C for 24 hours under static conditions. The geometric concentrations of the EOs’ stock solutions were then prepared in TSB or IVWM medium at concentrations ranging from 5 % to 0.002% (v/v) for T-EO and from 10% to 0.005% (v/v) for R-EO. After the biofilm’s incubation, the medium was gently removed, and 200 µL of EOs emulsions was added to the wells. The plates were incubated at 37°C for 24 hours under static conditions. Subsequently, the medium was removed, and 200 µL of 0.1% (w/v) TTC solution in the medium was added to each well for 2 hours at 37°C. Then MBEC values were assessed in the first colorless well. Subsequently, the medium was removed, and 200 µL of a solution of methanol (Stanlab, Poland) and acetic acid (Stanlab, Poland) in a 9:1 ratio was poured into the wells, and the plates were subjected to shaking (Mini-shaker PSU-2T, Biosan SIA, Latvia) for one hour at room temperature at 550 rpm. 100 µL of the solution was transferred to fresh 96-well plates (Wuxi Nest Biotechnology, China), and the absorbance of the solution was spectrophotometrically measured at 490 nm (Multiskan Go, Thermo Fisher Scientific, USA). Controls of bacterial growth (bacteria in TSB or IVWM medium) and medium sterility (medium only) were also prepared. To evaluate the percentage reduction of biofilm cells treated with EOs emulsions, the absorbance value of the sample was compared to the average bacterial growth absorbance.

### 2.9 Analysis of the Size of EOs Emulsion Droplets

The hydrodynamic diameters of the EOs droplets within emulsions were measured with the dynamic light scattering method. The analysis was performed using the Zetasizer Nano ZS ZEN3600 (Malvern Instruments Ltd., UK) equipped with a laser light source (λ=633 nm) and a detector operating at a scattering angle of 173°. Emulsions of EOs were prepared in TSB medium (Tryptic Soy Broth, Biomaxima, Poland) with 1% (v/v) Tween 20 (Pol-aura, Poland) and diluted in sterile purified water one thousand times prior to the measurement. The oil phase content was 5% (v/v) in the case of T-EO and 10% (v/v) regarding R-EO. The test was run at 25±0.1°C, and the samples were measured at least 5 times. The values of hydrodynamic diameters included in the work are Z-Average values.

### 2.10 Statistical analysis

Calculations were performed using Statistica software (Version 13; TIBCO Software Inc, Palo Alto, California, USA). The Hampel test was performed to identify outliers in the results of the biofilm biomass test, the biofilm metabolic activity assay, the biofilm thickness, and the share of live/ dead cells when all strains were analyzed together. The normality distribution and variance homogeneity were calculated with the Shapiro–Wilk and Levene tests, respectively. The t-test and the Mann-Whitney U test were performed to compare differences in biofilm biomass, biofilm metabolic activity, number of colony-forming units, thickness, and live/dead cell ratio between both media. The t-tests were used when normal distribution was determined (p>0.05). A Welch’s adjusted t-test was used for unequal variances (p<0.05). A Mann-Whitney U test was used when the normal distribution was not determined. The Pearson correlation was calculated to assess a linear correlation between biofilm biomass level and biofilm metabolic activity. Multivariate analysis of variance was performed to evaluate the effect of medium, strain, and EOs concentration on the reduction of biofilm cells after treatment with EOs. The results of statistical analyses with a significance level of p <0.05 were considered significant. The graphical abstract was created with BioRender.com (BioRender Inc, Switzerland, accessed on 17 May 2023)

## 3 Results

In the first line of the experiment, a GC-MS analysis was performed to evaluate the percentage composition of the EOs’ components. T-EO comprised 50.6% thymol, 19.2% p-cymene, and 9.1% γ-terpinene. Three main components of R-EO were: 21.1% α-pinene, 20.0% 1,8-cineole, and 18.5% camphor. A detailed list of the composition of the EOs is presented in **Supplementary Table 1**.

The ability of *S. aureus* strains to form biofilm biomass in standard TSB (Tryptic Soy Broth) or IVWM (In Vitro Wound Milieu) medium was evaluated using the crystal violet method (CV). The biofilms’ metabolic activity was assessed using tetrazolium chloride (TTC) staining **(Figure 1).**

**Figure 1.**
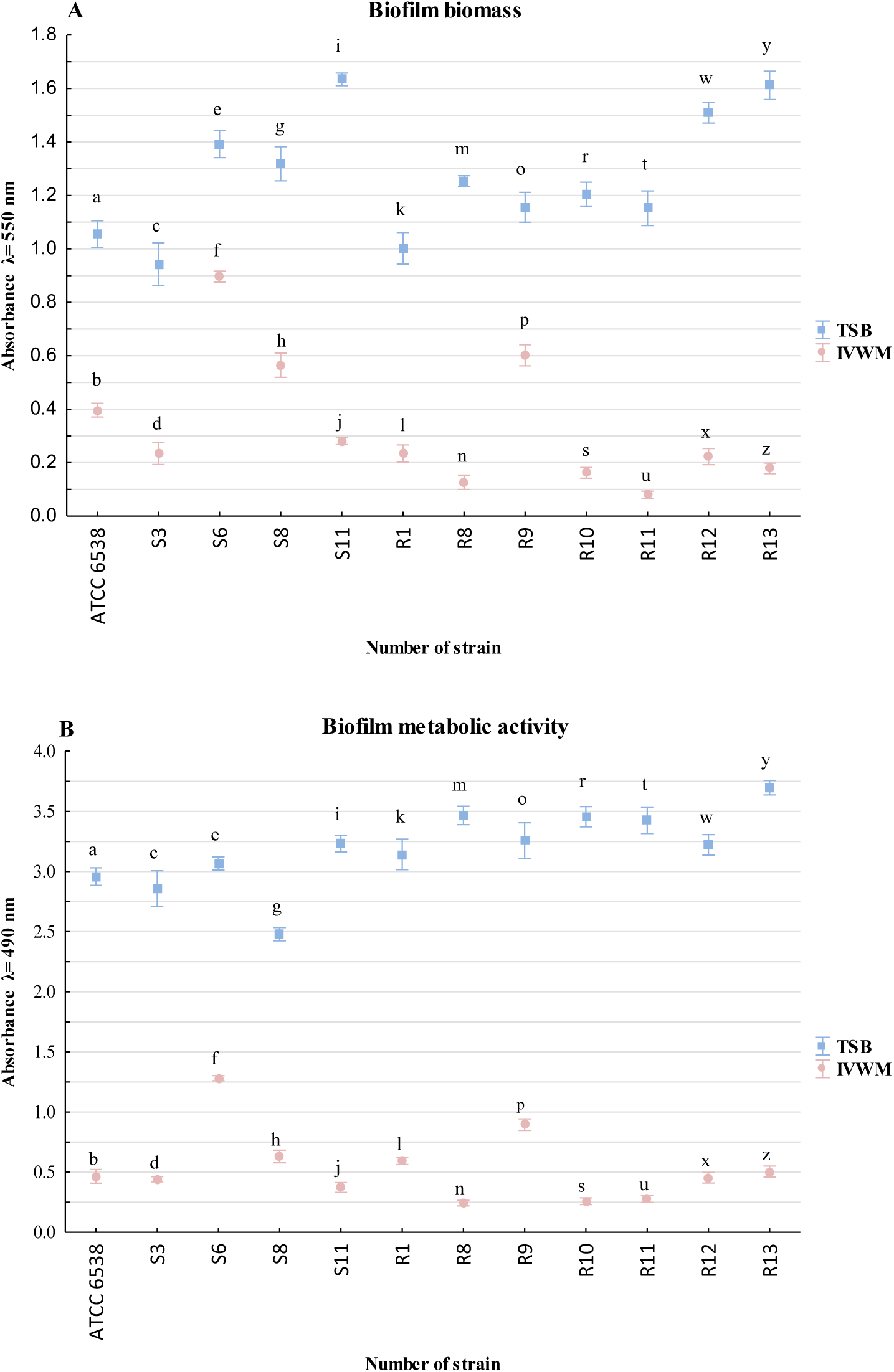
The average biofilm biomass **(A)** and metabolic activity **(B)** in Tryptic Soy Broth (TSB) or In Vitro Wound Milieu (IVWM) of analyzed reference (ATCC 6538, American Type Culture Collection) and clinical (S3, S6, S8, S11, R1, R8-R13) strains of *S. aureus*. The pairs of letters (a/b, c/d, e/f, g/h, i/j, k/l, m/n, o/p, r/s, t/u, w/x, y/z) refer to the statistically significant differences (p <0.05, t-test or Welch’s t-test for biofilm biomass assay, t-test or Welch’s t-test or Mann-Whitney U test for the biofilm metabolic activity assay). The error lines represent the standard error of the mean (n=12).

All strains were able to form a biofilm in TSB or IVWM medium. The biomass and metabolic activity of all biofilms were higher in TSB than in IVWM medium. For all strains, this trend was statistically significant (p <0.05, t-test or Welch’s t-test for the biofilm biomass assay, t-test or Welch’s t-test or Mann-Whitney U test for the biofilm metabolic activity assay, **Supplementary Tables 3, 4, Supplementary Figures 5, 6**), and it also remained statistically significant when the mean of all strains was calculated (p<0.05, Mann-Whitney U test), **Supplementary Figure 7, Supplementary Table 5)**. A significant linear correlation (p=0.001) between the level of biofilm biomass and metabolic activity for biofilms cultured in IVWM medium, but not in TSB medium (p=0.315), was observed (**Supplementary Table 6, Supplementary Figure 8**). The correlation coefficient was determined as moderate or very strong for TSB (r=0.32) or IVWM (r=0.83), respectively. To confirm data obtained by semiquantitative CV and TTC methods, the number of biofilm-forming cells of reference strain cultivated in TSB or IVWM was quantitatively cultured **(Table 2)**. The average number of biofilm-forming cells was significantly higher (p<0.05, t-test, **Supplementary Table 7, Supplementary Figure 9 A, B**) in TSB than IVWM medium. A macroscopic visualization of representative biofilm stained with crystal violet or tetrazolium chloride confirming differences in the level of biofilm biomass and metabolic activity between bacteria cultured in TSB or IVWM is presented in **Figure 2**. The biofilm in the TSB was highly confluent, i.e., biofilm-forming cells covered essentially the entire surface of the plate’s wells **(Figure 2 A, C**). Biofilm cells cultured in IVWM formed cells aggregates unequally distributed on the well’s surface **(Figure 2 B, D)**.

**Figure 2.**
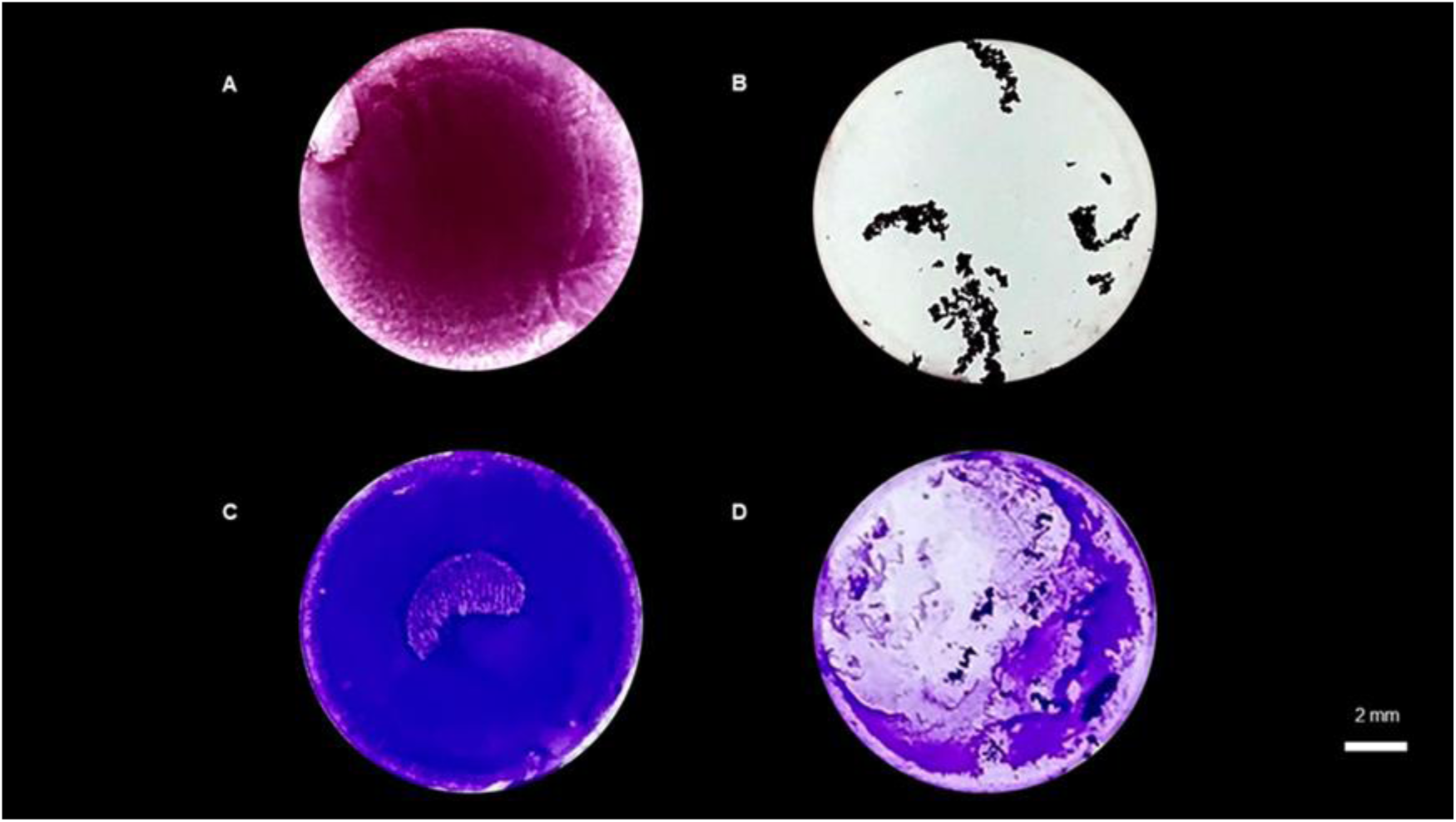
Macroscopic visualization of the *S. aureus* S11 biofilm formed on polystyrene. **A, B**-biofilm cultured in TSB or IVWM, respectively, and stained with tetrazolium chloride; **C, D**-biofilm cultured in TSB or IVWM, respectively, and stained with crystal violet. TSB-Tryptic Soy Broth, IVWM-In Vitro Wound Milieu. In macroscopic visualizations, the contrast was enhanced using GIMP software (Version 2.20.22, www.gimp.org, assessed 26.11.2020, original images are presented in **Supplementary Figure 1**). The scale bar is 2 mm.

**Table 2.** A mean number of Colony-Forming Units/mL (CFU/mL) of biofilm of *S. aureus* ATCC 6538 reference strain (American Type Culture Collection) cultured in TSB (Tryptic Soy Broth) or the IVWM (In Vitro Wound Milieu). SEM-standard error of the mean. A statistically significant difference (p<0.05, t-test) is marked with the pair of letters a/b.

| Number of CFU/mL |  |  |  |  |
| --- | --- | --- | --- | --- |
| Strain | TSB |  | IVWM |  |
| ATCC 6538 | Mean | SEM | Mean | SEM |
|  | 8.98E+14<sup>a</sup> | 1.07E+14 | 1.46E+12<sup>b</sup> | 6.65E+11 |

Next, the analysis of live and dead cells within the Z-axis (thickness) of the 24-hour-old staphylococcal biofilm structure was performed using LIVE/DEAD (L/D) dyeing and confocal microscopy **(Figure 3).** The staphylococcal biofilm cultured in TSB resembled the structure of a dense lawn **(Figure 3 A, C, E).** The biofilm-forming cells evenly covered the whole field of vision. In turn, the biofilm cultivated in IVWM formed mushroom-like structures of diversified sizes **(Figure 3 B, D, F).** The IVWM biofilm contained regions of differential thickness and cell density, including areas not covered with cells (black regions). Structural differences between biofilms cultured in TSB or IVWM medium were also observed in microscopic visualizations performed with the Scanning Electron Microscope (SEM) **(Figure 4).**

**Figure 3.**
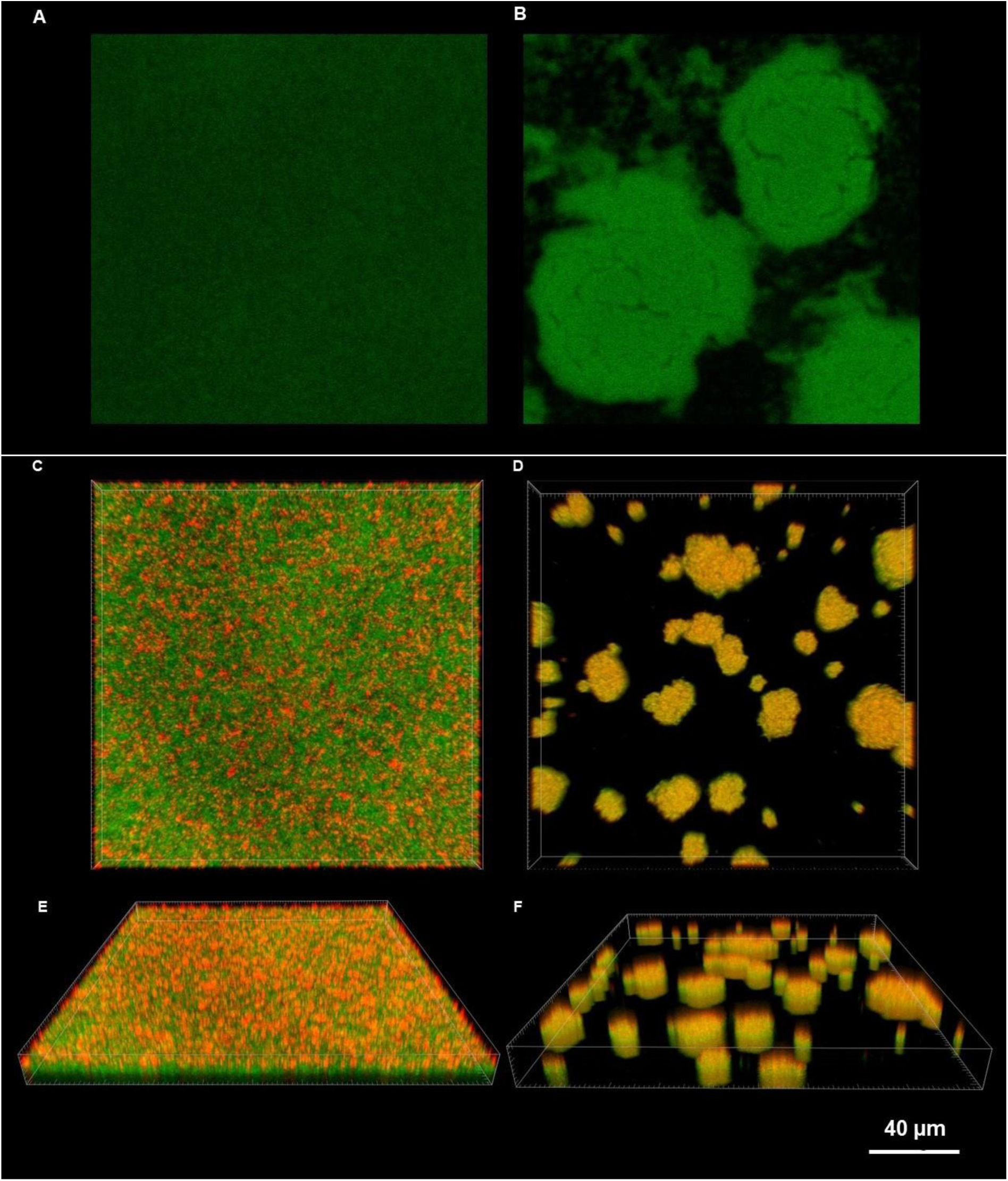
Microscopic visualizations of the *S. aureus* biofilm formed on polystyrene and stained with a LIVE/DEAD dye. **A, B**-an aerial perspective of the S8 biofilm cultured in TSB **(A)** or IVWM **(B)** – **C, D**-an aerial perspective of the S11 biofilm cultured in TSB **(C)** or IVWM **(D)**; **E, F-** the Z-axis image stack visualizing the S11 biofilm cultured in TSB **(E)** or IVWM **(F)** from the side aerial perspective. The red/orange color indicates staphylococcal cells of altered/damaged cell walls, while green-colored shapes show unaltered cell walls. TSB-Tryptic Soy Broth, IVWM-In Vitro Wound Milieu. The scale bar is 40µm. The confocal microscope SP8, magnification 25x.

**Figure 4.**
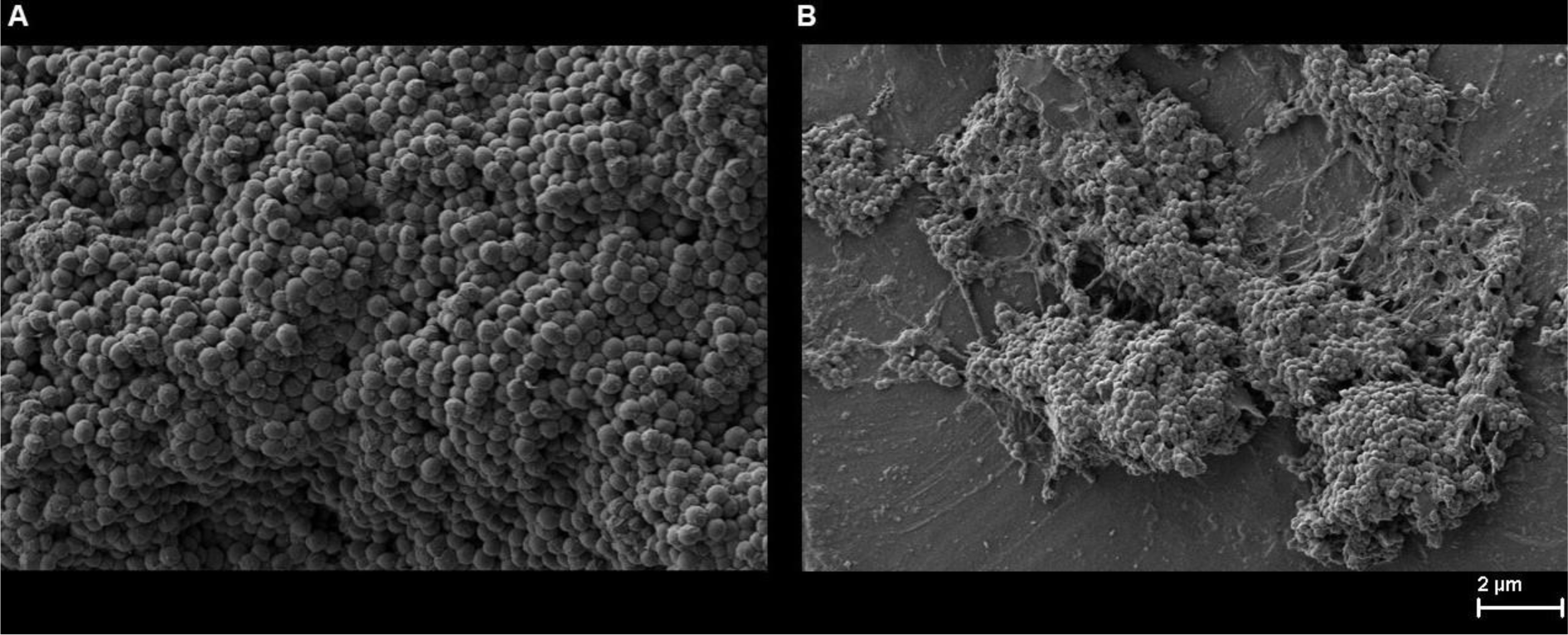
Microscopic visualizations of the *S. aureus* ATCC 6538 biofilm formed on agar and cultured in TSB **(A)** or IVWM **(B)** medium. TSB-Tryptic Soy Broth, IVWM-In Vitro Wound Milieu. The scale bar is 2µm. Scanning Electron Microscope Zeiss Auriga 60 (magnification 10000x).

The average thickness of the biofilm formed by all *S. aureus* strains in TSB medium was significantly lower than in IVWM medium (p<0.05, Mann-Whitney U test, **Figure 5**, **Supplementary Table 7, Supplementary Figure 9 C, D**).

**Figure 5.**
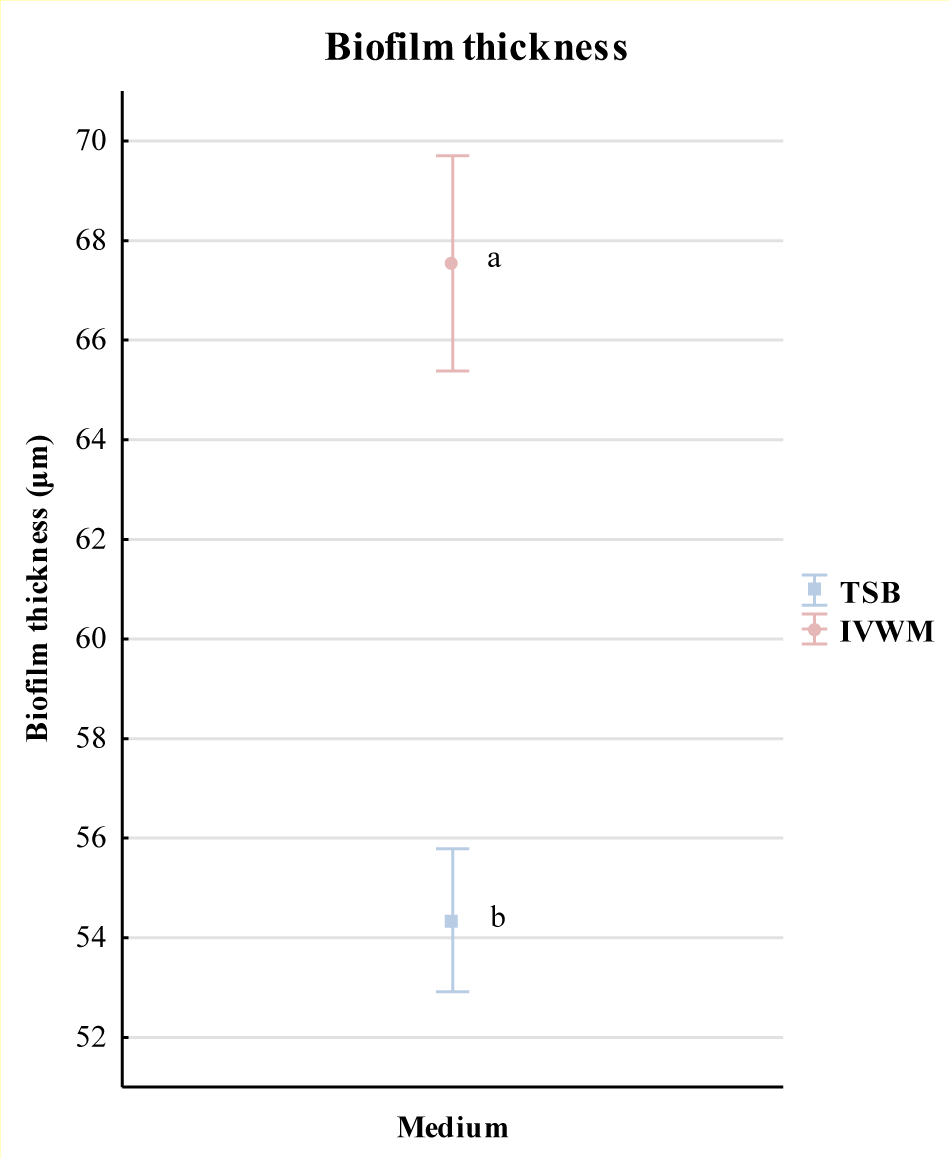
Comparison of the average thickness (µm) of all staphylococcal biofilms cultured in TSB (Tryptic Soy Broth) or IVWM (In Vitro Wound Milieu) medium measured with the confocal microscopy. The error lines represent the standard error of the mean (n=36). The statistically significant difference (p<0.05, Mann-Whitney U test) is marked with a pair of letters a/b.

In turn, the share of live cells was higher for ten (83% of strains) strains cultured in TSB medium than in IVWM **(Table 3)**. The average share of live cells was also significantly higher (p<0.05, Mann-Whitney U test) in TSB than in IVWM when all strains were analyzed together **(Supplementary Table 7, Supplementary Figure 9 E, F, G**).

**Table 3.** Average percentage (%) share of live and dead cells in biofilms of staphylococcal reference (ATCC 6538, American Type Culture Collection) or clinical (S3, S6, S8, S11, R1, R8-R13) strains cultured in TSB (Tryptic Soy Broth) or IVWM (In Vitro Wound Milieu) medium measured with the confocal microscope. SEM-standard error of the mean (n=3).

| Percentage (%) share of cells in *S. aureus* biofilms |  |  |  |  |  |  |
| --- | --- | --- | --- | --- | --- | --- |
| Strain number | TSB |  |  | IVWM |  |  |
|  | LIVE | DEAD | SEM | LIVE | DEAD | SEM |
| **ATCC 6538** | 84 | 16 | 0.1 | 80 | 20 | 1.0 |
| **S3** | 83 | 17 | 0.5 | 73 | 27 | 0.9 |
| **S6** | 76 | 24 | 2.3 | 61 | 39 | 3.1 |
| **S8** | 79 | 21 | 3.3 | 73 | 27 | 1.8 |
| **S11** | 74 | 26 | 2.4 | 63 | 37 | 1.5 |
| **R1** | 82 | 18 | 0.5 | 54 | 46 | 3.2 |
| **R8** | 86 | 14 | 1.7 | 67 | 33 | 0.5 |
| **R9** | 86 | 14 | 0.4 | 76 | 24 | 1.4 |
| **R10** | 66 | 34 | 1.1 | 68 | 32 | 0.9 |
| **R11** | 71 | 29 | 2.2 | 69 | 31 | 0.5 |
| **R12** | 70 | 30 | 0.5 | 74 | 26 | 1.3 |
| **R13** | 82 | 18 | 0.6 | 47 | 53 | 5.4 |

The analysis of the cellular spatial composition of the biofilms allowed us to distinguish their three sections: bottom one (B, the closest to the polystyrene surface), middle (M), and top (T, the farthest from the polystyrene surface) **(Figure 3**, **6, Supplementary Figure 2, Supplementary Table 2)**. Different patterns of live (green) and dead (red) share in parts (T, M, and B) of biofilms were revealed in each medium. Three (I, II, III) patterns were distinguished in TSB medium, and four (IV, V, VI, VII) in IVWM **(Figure 6**, **Supplementary Figure 2)**. The share of dead cells was higher in the T parts than in the M and B parts in 100% or 92% of staphylococcal biofilms grown in TSB or IVWM medium, respectively **(Figure 6**, **Supplementary Table 2)**. A higher share of live cells was observed in B (for 92% of strains) and M (for 83% of strains) parts of biofilms cultured in TSB medium than in IVWM **(Figure 6**, **Supplementary Figure 2, Supplementary Table 2)**. For 58% of staphylococcal biofilms, the share of live cells in T parts was higher in TSB medium than in IVWM.

**Figure 6.**
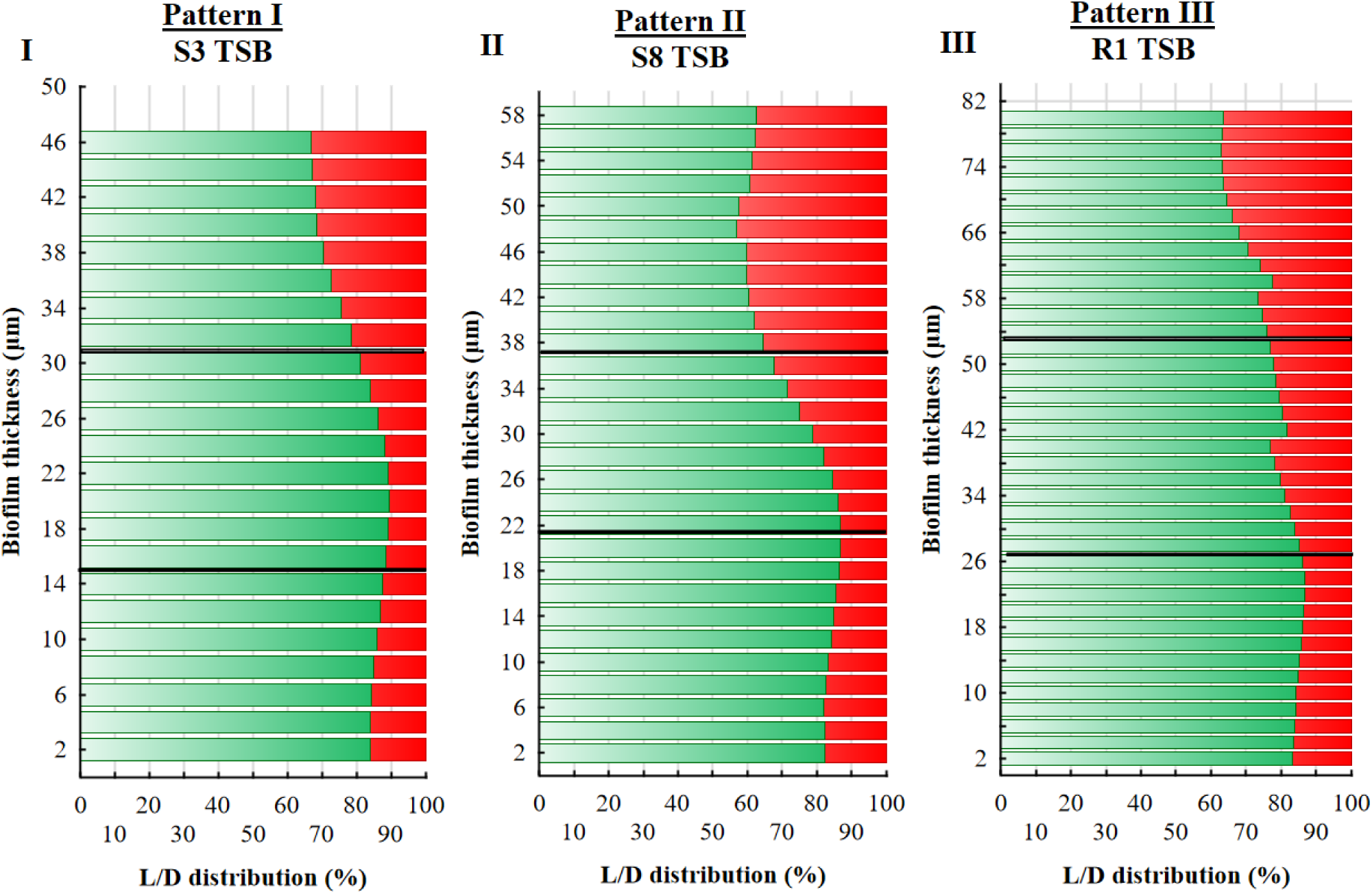
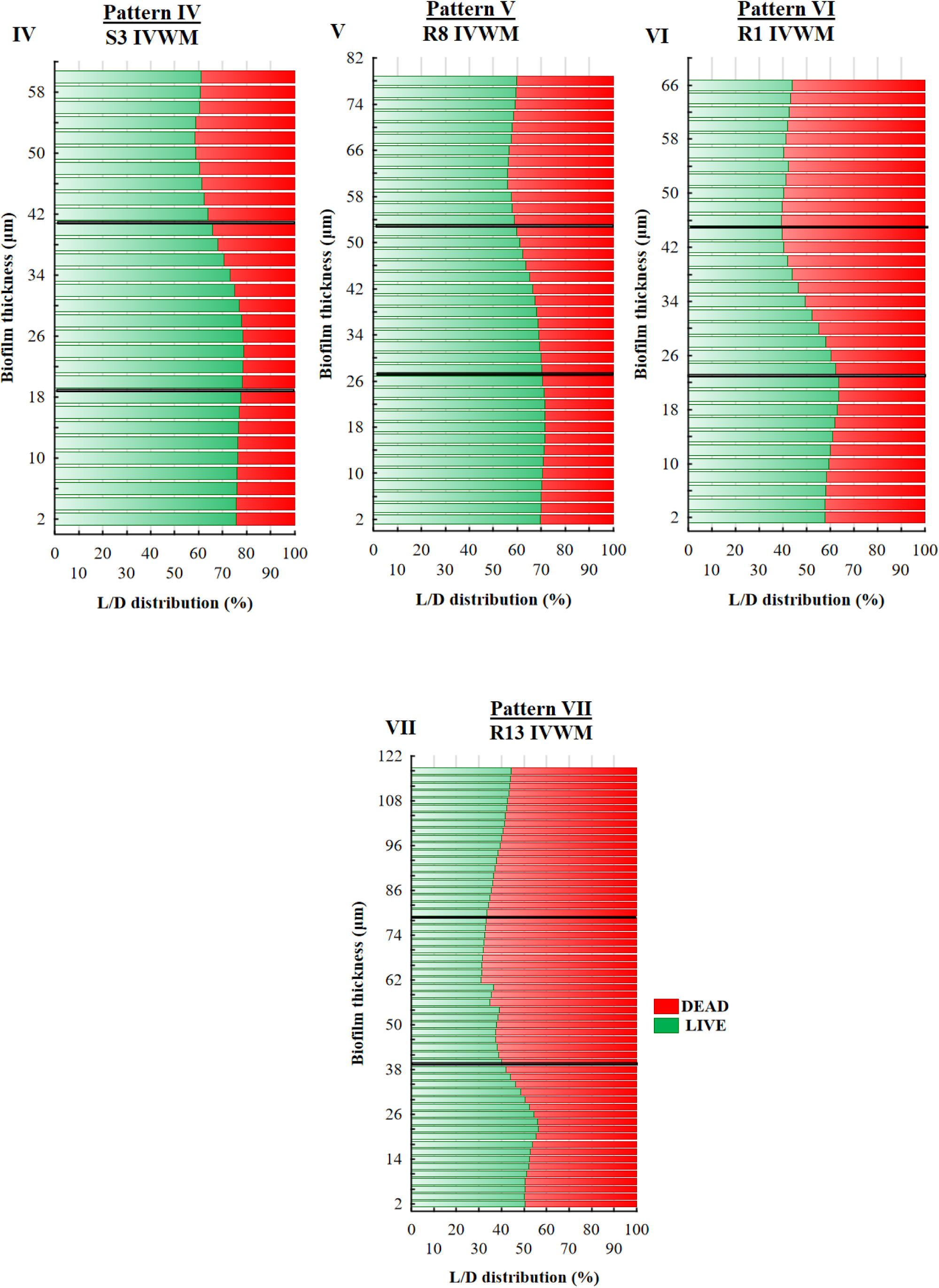
The main patterns of live (L, green) and dead (D, red) cells distribution (%) in staphylococcal biofilms across the Z-axis, cultured in TSB (Tryptic Soy Broth) or IVWM (In Vitro Wound Milieu) medium. The thickness of each section was 2 µm. Black lines divide graphs into the top (T), middle (M), and bottom (B) parts. **I, II**, and **III**-strains representing particular patterns of cells share in TSB; **IV, V, VI,** and **VII**-strains representing particular patterns of cells share in IVWM. S3, S8, R1, R8, R13-*S. aureus* clinical strains. Figures of the other strains are presented in **Supplementary Figure 2**. The confocal microscope SP8, magnification 25×.

The antimicrobial activity of the emulsified T- and R-EOs against planktonic *S. aureus* cells was evaluated using a microdilution technique **(Table 4)**. The applied emulsifier, Tween 20, did not affect the viability of staphylococcal cells (**Supplementary Figure 3).** The Minimal Inhibitory Concentrations (MIC) values of applied EOs were higher for 75% (T-EO) or 58% (R-EO) of strains cultivated in IVWM compared to strains cultivated in TSB medium. T-EO acted 2 to 4 times weaker against this 75% of strains in IVWM than in TSB medium. For 50% of strains treated with R-EO, the MIC values were twice higher in IVWM than in TSB, and for 8%, this parameter was sixty-four times higher in IVWM than in TSB.

**Table 4.** Antimicrobial activity of the tested EOs’ emulsions against planktonic (MIC (%) (v/v)) and biofilm cells (MBEC (%) (v/v)) of reference (ATCC 6538, American Type Culture Collection) or clinical (S3, S6, S8, S11, R1, R8-R13) strains of *S. aureus*. N/R indicates EOs where MBEC values were not reached at the highest applied concentration (10% (v/v)) of R-EO. Each EO’s MIC and MBEC values were compared in both media and marked green in a medium where the difference was higher than one geometric dilution, blue where the difference was one geometric dilution, and yellow where the parameters were equal. T-EO—thyme oil, R-EO—rosemary oil, TSB-Tryptic Soy Broth, IVWM-In Vitro Wound Milieu, MIC-Minimal Inhibitory Concentration, MBEC-Minimal Biofilm Eradication Concentration.

|  | MIC (%) |  |  |  | MBEC (%) |  |  |  |
| --- | --- | --- | --- | --- | --- | --- | --- | --- |
|  | T-EO |  | R-EO |  | T-EO |  | R-EO |  |
| Strain number | TSB | IVWM | TSB | IVWM | TSB | IVWM | TSB | IVWM |
| ATCC 6538 | 0.02 | 0.04 | 1.25 | 1.25 | 0.63 | 0.63 | N/R | N/R |
| S3 | 0.16 | 0.31 | 1.25 | 2.5 | 0.63 | 0.63 | 2.5 | N/R |
| S6 | 0.31 | 0.31 | 1.25 | 2.5 | 0.63 | 0.63 | N/R | N/R |
| S8 | 0.16 | 0.31 | 2.5 | 2.5 | 0.63 | 0.63 | N/R | N/R |
| S11 | 0.08 | 0.16 | 1.25 | 1.25 | 0.63 | 0.63 | N/R | N/R |
| R1 | 0.31 | 0.31 | 1.25 | 2.5 | 1.25 | 0.63 | N/R | N/R |
| R8 | 0.16 | 0.31 | 0.04 | 2.5 | 1.25 | 2.5 | N/R | N/R |
| R9 | 0.16 | 0.31 | 1.25 | 1.25 | 1.25 | 0.63 | N/R | N/R |
| R10 | 0.16 | 0.16 | 1.25 | 2.5 | 0.63 | 0.63 | N/R | N/R |
| R11 | 0.16 | 0.31 | 1.25 | 2.5 | 0.63 | 1.25 | N/R | N/R |
| R12 | 0.04 | 0.16 | 2.5 | 2.5 | 1.25 | 0.63 | N/R | N/R |
| R13 | 0.08 | 0.31 | 1.25 | 2.5 | 1.25 | 2.5 | N/R | N/R |

In the case of biofilms, T-EO exhibited higher activity than R-EO regardless of the medium applied **(Table 4)**. The MBEC (Minimal Biofilm Eradication Concentration) values of T-EO were determined for all staphylococcal strains and ranged from 2.5% (v/v) to 0.63% (v/v). The MBEC value of T-EO differed in 50% of strains, depending on the medium applied (TSB or IVWM). For half of these strains, the MBEC value was higher in IVWM than in TSB medium. In the case of R-EO, no MBEC values were achieved, therefore, the assessment of the biofilm cells’ reduction (%) was performed in the subsequent analyses.

The level of biofilm cell reduction (%) was above 90% for all strains in TSB or IVWM medium after the treatment with T-EO at concentrations of 2.5% - 0.63% (v/v) (**Supplementary Figure 4 A**). At a T-EO concentration of 0.31%, (v/v) lower reduction was observed for nine of twelve strains cultured in TSB comparing to the IVWM medium (**Figure 7 A**). The three strains with different patterns of reduction were: ATCC 6538, S6, and R9. The reduction of biofilm cells treated with R-EO was reached for all strains at concentrations ranging from 10% (v/v) to 0.63% (v/v) when cultured in TSB medium or from 10% (v/v) to 1.25% (v/v) in IVWM medium (**Figure 7 B, Supplementary Figure 4 B-F**). In concentration ranges of 10% (v/v) to 2.5% (v/v), a higher level of biofilm reduction was evaluated only for 30% of strains (R8-R11) cultured in IVWM medium than in TSB. The percentage reductions of staphylococcal biofilms after treatment with selected EOs concentrations are presented in **Figure 7** and **Supplementary Figure 4**. Multivariate analysis of variance was performed to evaluate the effect of medium, strain and EOs concentrations on the reduction of biofilm cells after treatment with T-EO or R-EO (**Supplementary Tables 8, 9, Supplementary Figures 10, 11**). In the case of both EOs, all of the factors significantly influenced the biofilm cells reduction. Additionally, for T-EO, there were significant interactions between each factor and all factors together. The biofilm cells reduction was significantly lower in TSB than in IVWM medium for T-EO and significantly higher in TSB than in IVWM for R-RO. In the case of R-EO, significant interactions occurred between medium and strain, strain and the oil concentration, and all three factors together.

**Figure 7.**
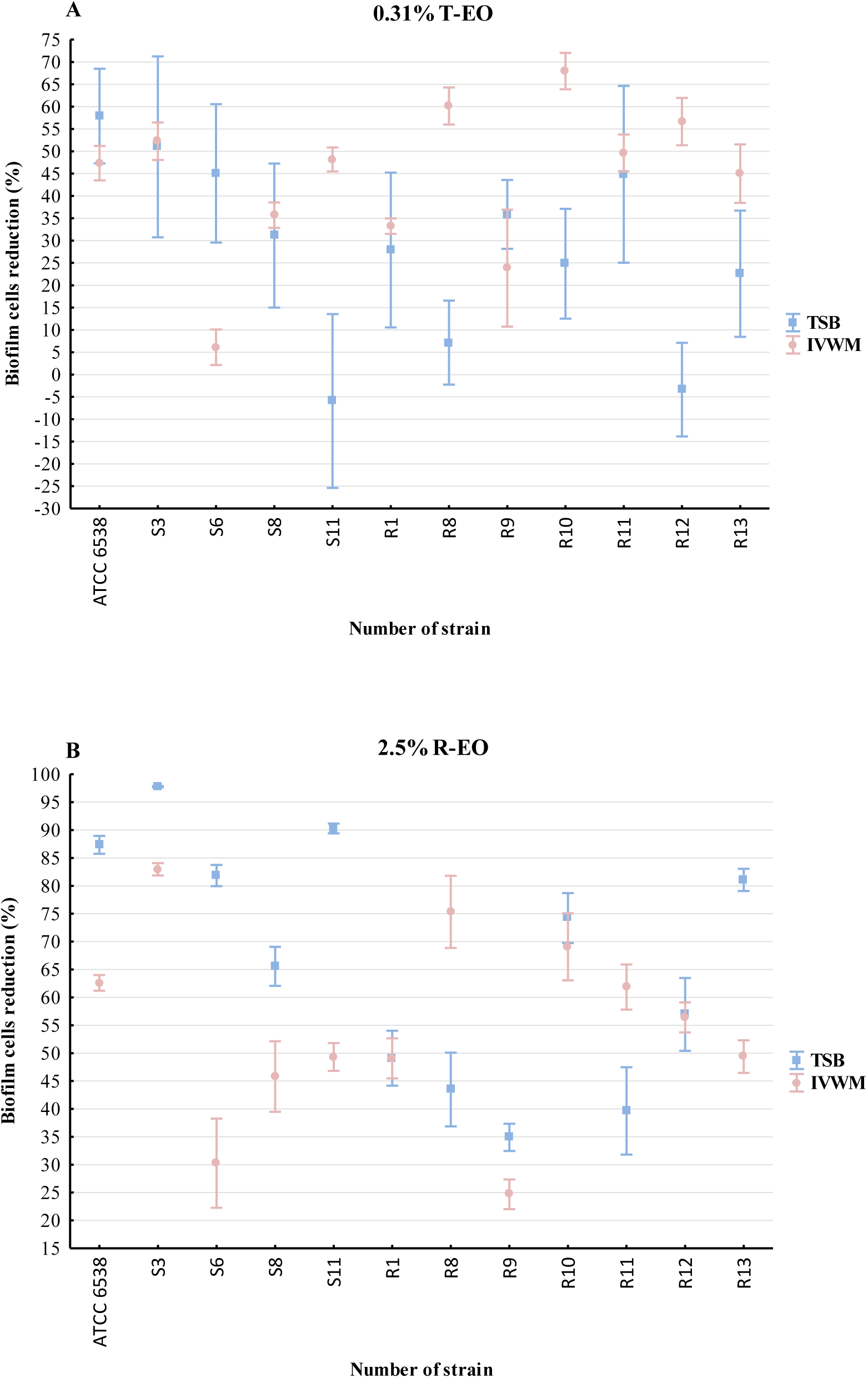
Average reduction (%) of biofilm cells of reference (ATCC 6538, American Type Culture Collection) or clinical (S3, S6, S8, S11, R1, R8-R13) strains of *S. aureus* after treatment with selected concentrations (%) (v/v) of thyme oil (**A**, T-EO) and rosemary oil (**B**, R-EO). The error lines represent the standard error of the mean (n=6). TSB-Tryptic Soy Broth, IVWM-In Vitro Wound Milieu.

Finally, the average diameters of the EOs droplets within the prepared formulations were measured. The droplets of T-EO and R-EO were 209±22 nm and 995±341 nm, respectively. The polydispersity index (PDI) of T-EO and R-EO’s emulsions was 0.46 and 0.68, respectively.

## 4 Discussion

Due to the limited usability of antibiotics in non-healing wound management and the emerging microbial resistance to antiseptics, there is a need to develop novel strategies for treating biofilm-related infections, such as utilizing natural compounds, e.g., essential oils (EOs). However, most *in vitro* research evaluates EOs’ antimicrobial activity against planktonic forms of microorganisms and in standard microbiological media. In contrast, the influence of EOs on microbial biofilm in a wound’s exudate-like medium should be assessed to accurately gauge their effectiveness at the infection site. Therefore, in this study, we applied a novel medium (In Vitro Wound Milieu, IVWM) containing serum, cell-matrix elements, and host factors that mimic the wound environment (Kadam et al., 2021). The biofilm characteristics and eradication following EO treatment were assessed in IVWM and a standard microbiological medium (Tryptic Soy Broth, TSB). Our first line of research involved the use of crystal violet and TTC (2,3,5-triphenyl-tetrazolium chloride) staining to evaluate the strains’ ability to form a biofilm (**Figure 1**). The results revealed high intraspecies variability in biofilm biomass and metabolic activity, regardless of the medium used.

The biomass, metabolic activity, and cell number of *S. aureus* biofilms were higher in TSB than in IVWM medium (**Figures 1**, **2**). Unlike TSB, which primarily contains proteins, glucose, and sodium chloride, IVWM presents host factors that may impair biofilm formation (**Table 1**). Studies indicate that lactoferrin inhibits bacterial growth by binding iron, which restricts its availability for bacteria, or through direct interaction with negatively charged regions of bacterial membranes, causing cell damage (Aguila et al., 2001; Ammons and Copié, 2013). Lactoferrin can also obstruct biofilm formation by preventing its adhesion, disrupting existing structures, and significantly altering the expression of genes responsible for cell metabolism (Roseanu et al., 2010; Ammons and Copié, 2013). According to Abraham et al., non-protein compounds of bovine serum (which constitutes 70% of IVWM medium) inhibit staphylococcal biofilm formation (Abraham and Jefferson, 2010).

Moreover, the biofilm in TSB was highly confluent (**Figure 2 A, C**), while the biofilm cultured in IVWM formed cellular aggregates unevenly distributed on the well’s surface. However, these were covered with an extracellular matrix to a greater extent than the biofilms formed in the TSB (**Figure 2 B, D**). These findings confirm that IVWM medium accurately reflects biofilm structure under *in vivo* conditions because bacterial aggregates were also observed in biopsy materials from chronic wounds (Kirketerp-Møller et al., 2008; Fazli et al., 2009; Bay et al., 2018). *S. aureus* is equipped with surface proteins that bind to fibrinogen and fibronectin, leading to the formation of cell clusters. Aggregation facilitates bacterial evasion from host’s immune system, including phagocytosis (Crosby et al., 2016; Pestrak et al., 2020). Nevertheless, cell aggregation has been reported to limit bacterial attachment to surfaces such as steel or hydroxyapatite. This may be explained by the fact that larger particle sizes lead to higher drag forces and lower attachment (Pestrak et al., 2020). This phenomenon presumably accounts for the significantly lower biofilm mass formed in IVWM than in TSB, demonstrated in our study.

A positive high correlation (r=0.83) was observed between the level of biofilm biomass and metabolic activity for biofilms cultured in IVWM medium but not in TSB medium (**Supplementary Table 6, Supplementary Figure 8**). One explanation for the lack of aforementioned correlation in TSB could be that biofilm structure is metabolically differentiated, and regions with decreased metabolic activity may be present in biofilms where an abundant extracellular matrix hinders the diffusion of nutrients and oxygen (Marcos-Zambrano et al., 2014; Xu et al., 2016). To extensively assess biofilm characteristics in both media, we analyzed the biofilm’s 3D structure using LIVE/DEAD staining and confocal microscopy. The biofilm cells in TSB are evenly distributed, thus forming a thinner structure than in IVWM, where the cells are clustered in aggregates with more cell layers, forming mushroom-like structures. However, in IVWM, there are also cell-free areas between these aggregates (**Figures 3**, **4**). The different spatial distribution of cells in both media may be the cause of the lower thickness of the biofilm formed in the TSB medium than in the IVWM medium (**Figure 5**). It was indicated that the percentage of live cells was higher in biofilms cultured in TSB medium than in IVWM (**Table 3**, **Supplementary Figure 89G**). The finding aligns with the results of the metabolic analysis assay, where the cells’ metabolic activity was higher in TSB than in IVWM medium (**Figure 1B**). This may be due to less favorable conditions for biofilm formation in IVWM, such as the presence of antibiofilm compounds and a lower concentration of nutrients. When analyzing the cellular spatial composition of the biofilms, we demonstrated that in both media, the share of dead cells was higher in the top (T) than in the middle (M) and bottom (B) parts (**Figure 6**, **Supplementary Figure 2, Supplementary Table 2**). In IVWM medium, this could be a biofilm protection mechanism against antimicrobial substances. Dead cells are preferentially localized and act as a barrier, limiting the diffusion of antimicrobials into deeper layers of the biofilm (Chambless et al., 2006). Subsequently, *S. aureus* planktonic and biofilm cells were cultured in both media and treated with T- and R-EO. EOs showed significant antistaphylococcal activity, though intraspecies variability was observed (**Table 4**, **Figure 7**, **Supplementary Figure 4**). This is consistent with the results presented by our research team and other researchers (Tohidpour et al., 2010; Kot et al., 2018; Brożyna et al., 2021) and indicates the necessity of including a broad spectrum of various strains in the tests. Differences in EOs’ MIC and MBEC values were observed between experimental settings using IVWM or TSB medium, and they were higher in the case of planktonic forms than in biofilms. EOs exhibited lower antimicrobial activity against planktonic forms in IVWM than in TSB medium for more than half of the strains. This may be due to the impairment of EOs activity by bovine serum albumin (Juven et al., 1994; Hammer and Carson, 2010). EOs influence bacteria mainly by binding to the cell wall/membrane and leading to the disruption of its integrity. Albumin presumably binds to the hydrophobic components of EOs and hinders the interaction of EOs with bacterial membrane proteins, which decreases their efficacy (Juven et al., 1994; Veldhuizen et al., 2007). In general, a lower level of biofilm reduction was obtained after treatment with R-EO in IVWM than in TSB medium (**Figure 7**, **Supplementary Figure 4**). Regarding the antibiofilm activity of T-EO, higher cell reduction was achieved for 75% of strains cultured in IVWM than in TSB medium.

Previous studies demonstrated that clustered bacterial cells (like those in IVWM medium) exhibit increased tolerance to antimicrobial substances (Pestrak et al., 2020). Thicker biofilms, observed in IVWM medium, are also more difficult to eradicate because antimicrobial agents’ penetration through the biofilm structure is reduced. The higher antibiofilm activity of T-EO in IVWM than in TSB medium against specific strains suggests that the EO influences biofilm not only by direct interaction with the cell wall but also through other mechanisms, which are enhanced with IVWM medium components. In both the present study and our previous research, we demonstrated that key biofilm characteristics differ significantly depending on the medium in which microorganisms are cultivated (Paleczny et al., 2021, 2022). These differences translate to variances in the results of the antimicrobial activity of the tested substances. Therefore, using only standard microbiological media in *in vitro* studies to evaluate the antimicrobial substances’ efficacy may lead to over- or underestimation of their effect.

Although intraspecies variability was observed, we demonstrated the high effectiveness of EOs in the eradication of *S. aureus* biofilm and planktonic forms in both TSB and IVWM media. EOs, especially T-EO, are promising agents for the treatment of biofilm-related wound infections since, in specific cases, their activity in the IVWM medium was higher than in TSB.

The research results underscore the importance of using a medium that reflects the state of the infection site (e.g., wound exudate) and including a high number of tested strains in *in vitro* studies. This understanding will help develop effective treatments for biofilm-related infections and better manage wound healing processes in the future.

## Highlights

- Biofilm cultured in IVWM is thicker, and the biomass, metabolic activity, and cell number are higher than in TSB medium.
- T- and R-EO exhibited lower antimicrobial activity against planktonic forms in IVWM than in TSB medium.
- A lower level of biofilm reduction was obtained after treatment with R-EO in IVWM than in TSB medium.
- A higher level of biofilm reduction was obtained after treatment with T-EO in IVWM than in TSB medium.

## Conflict of Interest

The authors declare that the research was conducted in the absence of any commercial or financial relationships that could be interpreted as a potential conflict of interest.

## Author Contributions

MB, AJ designed the research. MB, JP, WK, KM, and AJ performed the experiments. MB and JD performed the statistical analysis. MB wrote the first draft of the manuscript. MB, JD, and AJ analyzed the data and prepared graphics. WK and KM wrote sections of the manuscript. AJ and KFM revised and edited the manuscript. JD and AJ revised the English. AJ, JD supervised the work. All authors contributed to the article and approved the submitted version.

## Funding

This research was funded in whole by the National Science Centre, Poland (Grant No. 2021/41/N/NZ6/03305). For the purpose of Open Access, the author has applied a CC-BY public copyright license to any Author Accepted Manuscript (AAM) version arising from this submission.

## Notes

### Competing Interest Statement

The authors have declared no competing interest.

