## Supplement for "Chronic Wound Milieu Challenges Essential Oils’ Antibiofilm Activity"

Adam Junka*

**Supplementary Table 1**. Ingredients of T-EO **(A)** and R-EO **(B)** measured with GC-MS (Gas Chromatography-Mass Spectrometry). Components in line with Polish Pharmakopea XI standards are marked green; the ones not in line are marked red. RI - retention index, RT – retention time, T-EO- thyme essential oil, R-RO- rosemary essential oil.

| **(A) T-EO** | | | |
|---|---|---|---|
| **RI** | **RT** | **Compound** | **T-EO** |
| 925 | 8.94 | α-thujene | 0.38 |
| 932 | 9.19 | α-pinene | 0.91 |
| 948 | 9.77 | camphene | 1.06 |
| 976 | 10.75 | β-pinene | 0.31 |
| 988 | 11.19 | myrcene | 1.11 |
| 1016 | 12.19 | α-terpinene | 1.81 |
| **1024** | **12.50** | **p-cymene** | **19.20** |
| 1028 | 12.65 | limonene | 0.37 |
| 1030 | 12.70 | eucalyptol | 0.28 |

| | | | |
|---|---|---|---|
| **1058** | **13.74** | **γ-terpinene** | **9.06** |
| 1099 | 15.28 | linalool | 3.21 |
| 1147 | 17.01 | camphor | 0.62 |
| 1172 | 17.92 | borneol | 1.98 |
| 1180 | 18.23 | terpinen-4-ol | 1.02 |
| 1195 | 18.76 | α-terpineol | 0.42 |
| 1238 | 20.27 | thymol methyl ether | 0.46 |
| **1291** | **22.15** | **thymol** | **50.59** |
| 1297 | 22.36 | carvacrol | 5.65 |
| 1426 | 26.37 | caryophyllene | 1.66 |

Polish Pharmakopea XI ranges for thyme oil, thymol chemotype:
α- thujene: 0.2 percent to 1.5 percent
β-myrcene: 1.0 percent to 3.0 percent
α-terpinene: 0.9 percent to 2.6 percent
p-cymene: 14.0 percent to 28.0 percent
γ-terpinene: 4.0 percent to 12.0 percent
linalool: 1.5 percent to 6.5 percent
terpinen-4-ol: 0.1 percent to 2.5 percent
carvacrol methyl ether: 0.05 percent to 1.5 percent
thymol: 37.0 percent to 55.0 percent
carvacrol: 0.5 percent to 5.5 percent

| **(B) R-EO** | | | |
|---|---|---|---|
| **RI** | **RT** | **Compound** | **R-EO** |
| 921 | 8.81 | tricyclene | 0.41 |
| 925 | 8.94 | α-thujene | 0.11 |
| **933** | **9.22** | **α-pinene** | **21.07** |
| 948 | 9.77 | camphene | 8.81 |
| 952 | 9.90 | thuja-2,4(10)-diene | 0.30 |
| 976 | 10.76 | β-pinene | 3.78 |
| 984 | 11.04 | octen-2-ol | 0.20 |
| 989 | 11.20 | myrcene | 2.70 |
| 1005 | 11.80 | α-phellandrene | 0.52 |
| 1008 | 11.89 | 3-carene | 0.61 |
| 1016 | 12.19 | α-terpinene | 0.65 |
| 1024 | 12.48 | p-cymene | 2.46 |
| 1029 | 12.66 | limonene | 3.88 |
| **1032** | **12.79** | **1,8-cineole** | **19.98** |
| 1035 | 12.91 | trans-β-ocimene | 0.16 |
| 1057 | 13.73 | γ-terpinene | 1.01 |
| 1084 | 14.73 | terpinolene | 0.68 |

| | | | |
|---|---|---|---|
| 1099 | 15.28 | linalool | 0.71 |
| **1148** | **17.05** | **camphor** | **18.52** |
| 1172 | 17.92 | borneol | 3.54 |
| 1180 | 18.23 | terpinen-4-ol | 0.68 |
| 1194 | 18.76 | α-terpineol | 2.32 |
| 1206 | 19.18 | verbenone | 1.91 |
| 1284 | 21.88 | bornyl acetate | 1.23 |
| 1369 | 24.74 | copaene | 0.15 |
| 1419 | 26.37 | β-caryophyllene | 2.76 |
| 1456 | 27.50 | α-humulene | 0.50 |
| 1479 | 28.23 | γ-muurolene | 0.11 |
| 1517 | 29.41 | δ-cadinene | 0.12 |

For rosemary oil, Spanish type**,** the percentages are within the following ranges:

α-pinene: 18 percent to 26 percent
camphene: 8.0 percent to 12.0 percent
β-pinene: 2.0 percent to 6.0 percent
β-myrcene: 1.5 percent to 5.0 percent
limonene: 2.5 percent to 5.0 percent
cineole: 16.0 percent to 25.0 percent
p-cymene: 1.0 percent to 2.2 percent
camphor: 13.0 percent to 21.0 percent
bornyl acetate: 0.5 percent to 2.5 percent
α-terpineol: 1.0 percent to 3.5 percent
borneol: 2.0 percent to 4.5 percent
verbenone: 0.7 percent to 2.5 percent

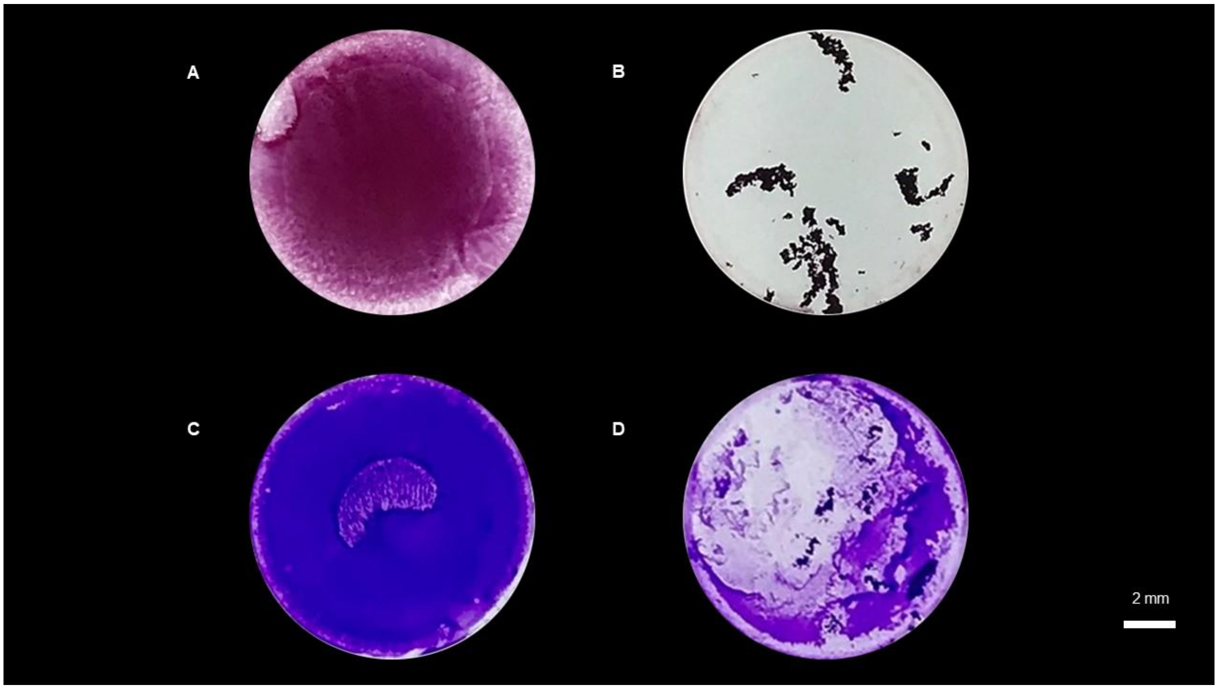


**Supplementary Figure 1**. Original images of *S. aureus* S11 biofilm formed on polystyrene. **A, B**- biofilm cultured in TSB (**A**) or IVWM (**B**) and stained with tetrazolium chloride; **C, D**- biofilm cultured in TSB (**C**) or IVWM (**D**) and stained with crystal violet. TSB- Tryptic Soy Broth, IVWM- In Vitro Wound Milieu. The scale bar is 2 mm.

**Supplementary Table 2.** Average percentage share of live and dead cells in biofilms of staphylococcal reference (ATCC 6538, American Type Culture Collection) or clinical (S3, S6, S8, S11, R1, R8-R13) strains cultured in TSB (Tryptic Soy Broth) or IVWM (In Vitro Wound Milieu) medium measured with a confocal microscope. Biofilms were divided into three parts along the Z-axis: top (T), middle (M), and bottom (B). The share of cells in each part was compared in both media and marked green in medium where the parameter was higher and red where it was lower.

| | | | **Share (%) of cells in *S. aureus* biofilms** | |
|---|---|---|---|---|
| **Strain number** | **Part of biofilm** | **Type of cells** | **TSB** | **IVWM** |
| **ATCC 6538** | T | Live | 68 | 71 |
| | | Dead | 32 | 29 |
| | M | Live | 84 | 79 |
| | | Dead | 16 | 21 |
| | B | Live | 87 | 82 |
| | | Dead | 13 | 18 |
| **S3** | T | Live | 71 | 61 |
| | | Dead | 29 | 39 |
| | M | Live | 87 | 74 |
| | | Dead | 13 | 26 |

| | | | | |
|---|---|---|---|---|
| | B | Live | 85 | 76 |
| | | Dead | 15 | 24 |
| **S6** | T | Live | 56 | 48 |
| | | Dead | 44 | 52 |
| | M | Live | 73 | 60 |
| | | Dead | 27 | 40 |
| | B | Live | 79 | 69 |
| | | Dead | 21 | 31 |
| **S8** | T | Live | 60 | 64 |
| | | Dead | 40 | 36 |
| | M | Live | 78 | 76 |
| | | Dead | 22 | 24 |
| | B | Live | 84 | 76 |
| | | Dead | 16 | 24 |
| **S11** | T | Live | 61 | 54 |
| | | Dead | 39 | 46 |
| | M | Live | 73 | 62 |
| | | Dead | 27 | 38 |
| | B | Live | 79 | 67 |
| | | Dead | 21 | 33 |
| **R1** | T | Live | 69 | 41 |
| | | Dead | 31 | 59 |
| | M | Live | 80 | 50 |
| | | Dead | 20 | 50 |
| | B | Live | 85 | 60 |
| | | Dead | 15 | 40 |
| **R8** | T | Live | 72 | 58 |
| | | Dead | 28 | 42 |
| | M | Live | 88 | 66 |
| | | Dead | 12 | 34 |
| | B | Live | 89 | 71 |
| | | Dead | 11 | 29 |
| **R9** | T | Live | 74 | 68 |
| | | Dead | 26 | 32 |
| | M | Live | 87 | 70 |
| | | Dead | 13 | 30 |
| | B | Live | 88 | 81 |
| | | Dead | 12 | 19 |
| **R10** | T | Live | 53 | 58 |
| | | Dead | 47 | 42 |
| | M | Live | 63 | 66 |
| | | Dead | 37 | 34 |
| | B | Live | 73 | 71 |
| | | Dead | 27 | 29 |
| **R11** | T | Live | 56 | 58 |
| | | Dead | 44 | 42 |

| | | | | |
|---|---|---|---|---|
| | M | Live | 67 | 63 |
| | | Dead | 33 | 37 |
| | B | Live | 77 | 74 |
| | | Dead | 23 | 26 |
| **R12** | T | Live | 51 | 65 |
| | | Dead | 49 | 35 |
| | M | Live | 70 | 74 |
| | | Dead | 30 | 26 |
| | B | Live | 75 | 77 |
| | | Dead | 25 | 23 |
| **R13** | T | Live | 65 | 39 |
| | | Dead | 35 | 61 |
| | M | Live | 82 | 35 |
| | | Dead | 18 | 65 |
| | B | Live | 86 | 51 |
| | | Dead | 14 | 49 |

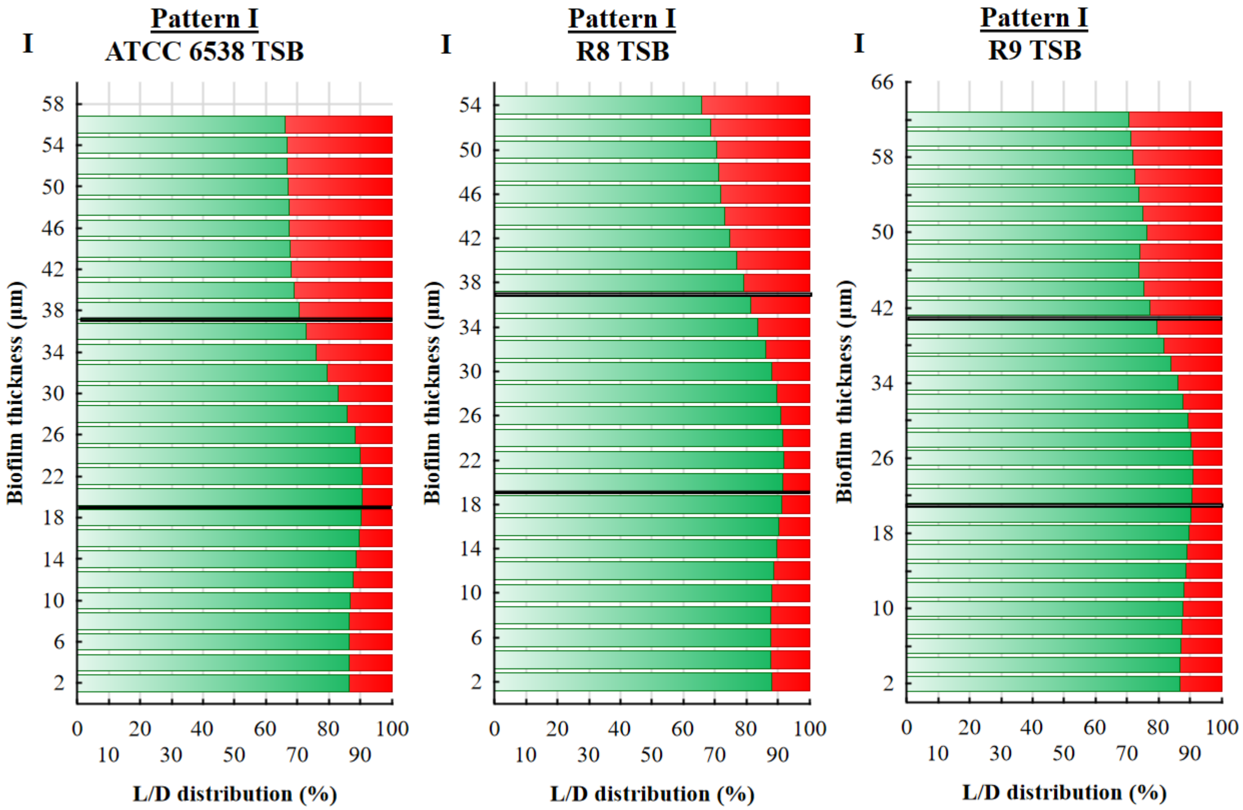

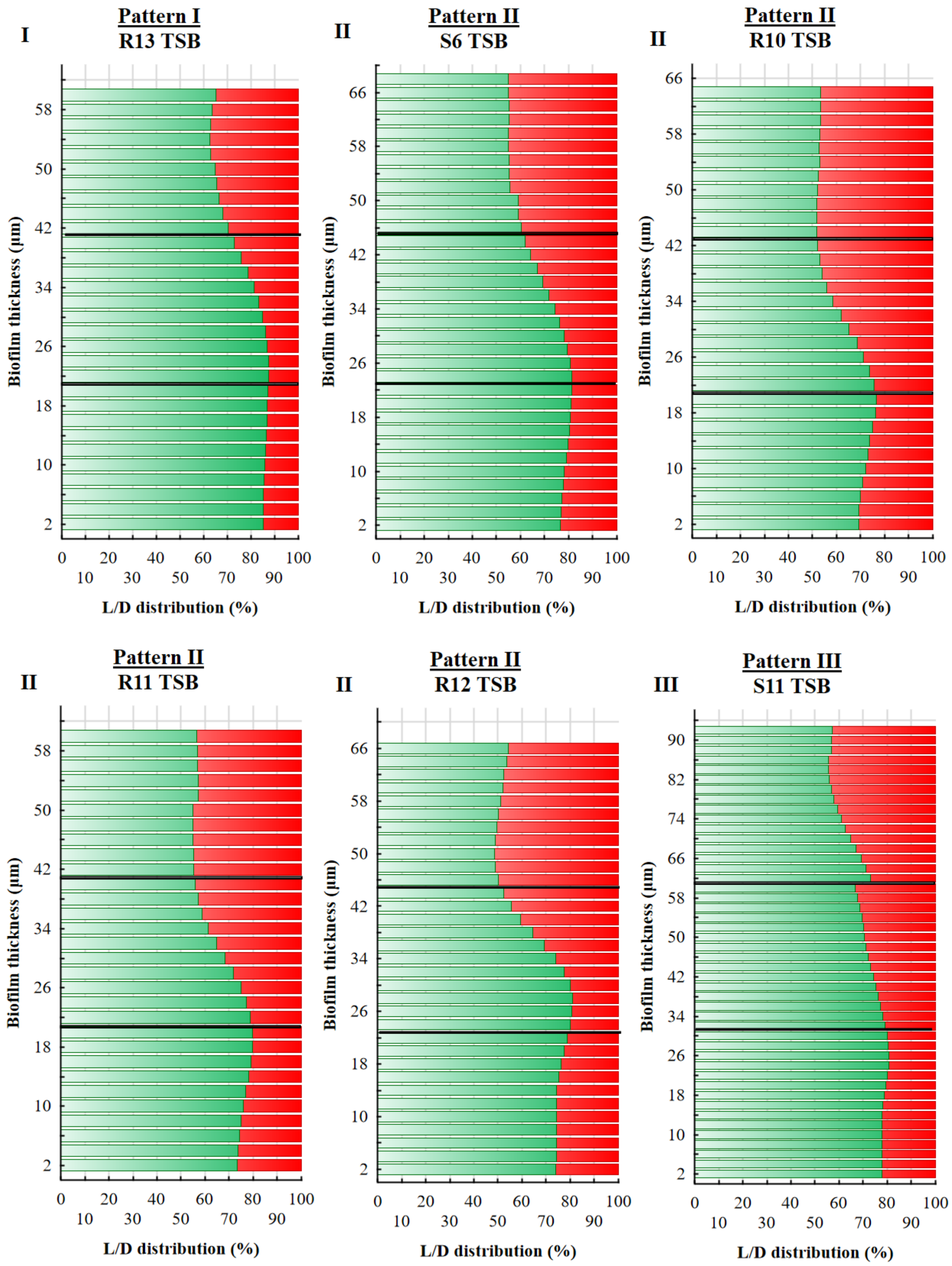

I
Pattern I
R13 TSB
II
Pattern II
S6 TSB
II
Pattern II
R10 TSB
II
Pattern II
R11 TSB
II
Pattern II
R12 TSB
III
Pattern III
S11 TSB
Biofilm thickness (µm)
L/D distribution (%)

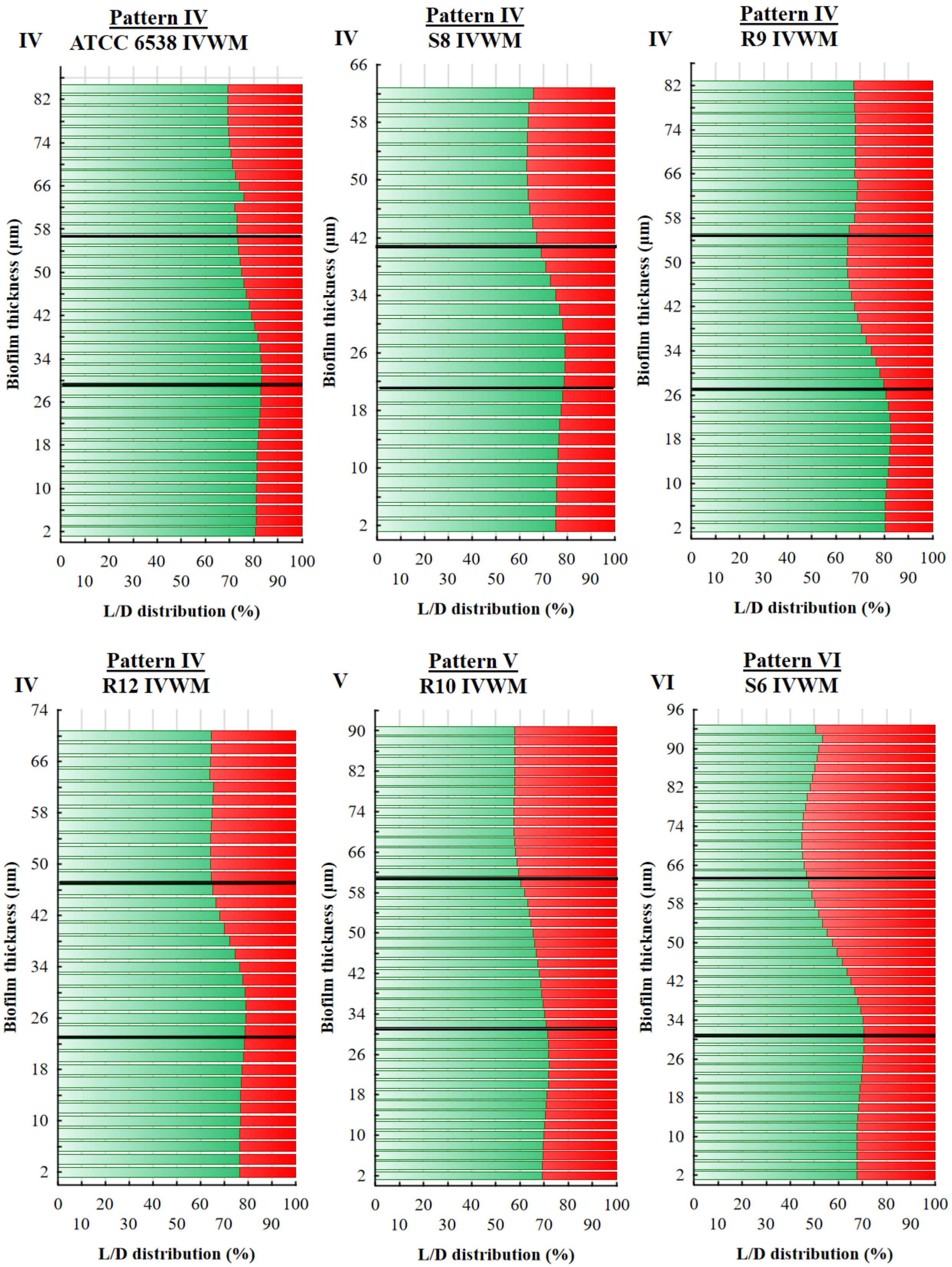
IV
Pattern IV
ATCC 6538 IVWM
Biofilm thickness (µm)
L/D distribution (%)
IV
Pattern IV
S8 IVWM
Biofilm thickness (µm)
L/D distribution (%)
IV
Pattern IV
R9 IVWM
Biofilm thickness (µm)
L/D distribution (%)
IV
Pattern IV
R12 IVWM
Biofilm thickness (µm)
L/D distribution (%)
V
Pattern V
R10 IVWM
Biofilm thickness (µm)
L/D distribution (%)
VI
Pattern VI
S6 IVWM
Biofilm thickness (µm)
L/D distribution (%)

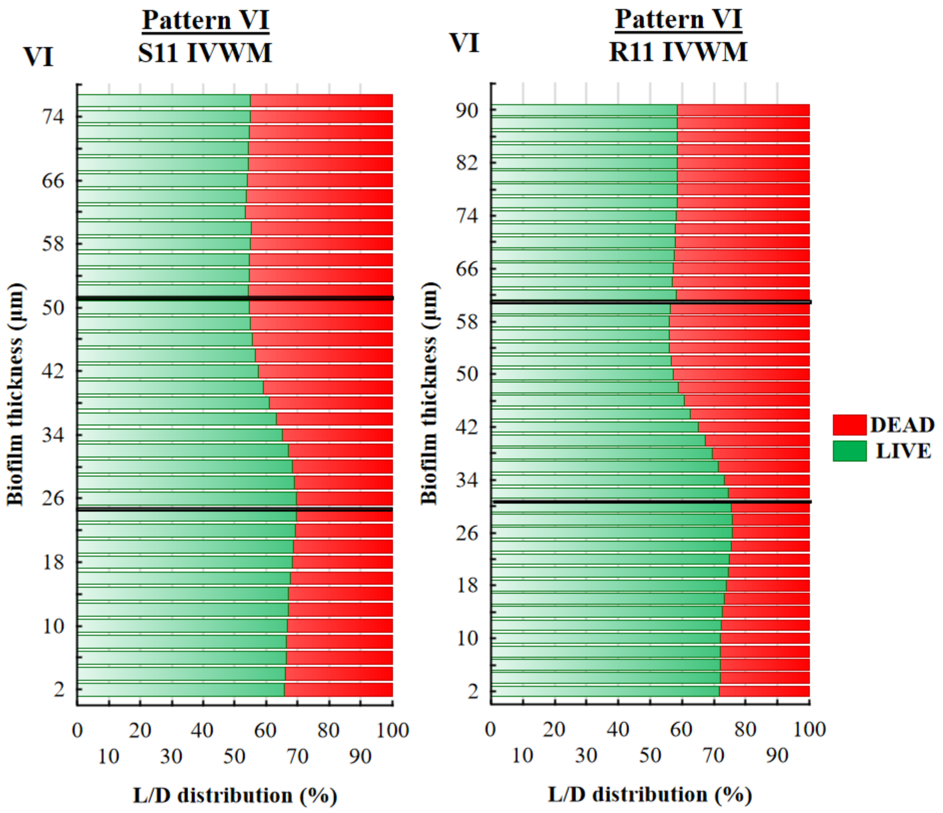


**Supplementary Figure 2.** The main patterns of live (L, green) and dead (D, red) cells distribution (%) in staphylococcal biofilms across the Z-axis, cultured in TSB (Tryptic Soy Broth) or IVWM (In Vitro Wound Milieu) medium. The thickness of each section was 2 μm. Black lines divide graphs into the top (T), middle (M), and bottom (B) parts. **I, II, III**- strains of particular patterns of cells share in TSB; **IV, V, VI**-strains of particular patterns of cells share in IVWM. ATCC 6538- *S. aureus* reference strain (American Type Culture Collection), S6, S8, S11, R8-R13- clinical strains of *S. aureus*. The confocal microscope SP8, magnification 25×.

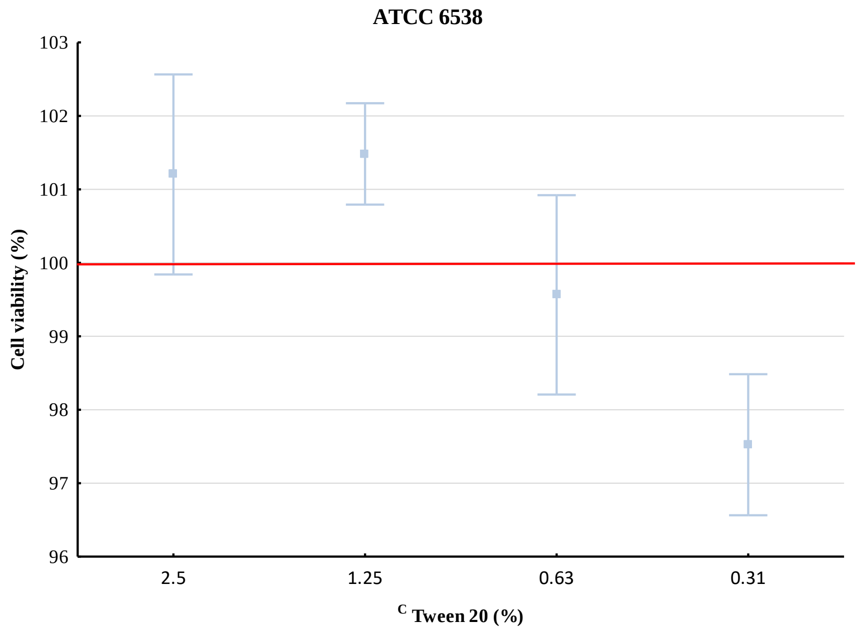

**Supplementary Figure 3.** Average viability (%) of *S. aureus* ATCC 6538 (American Type Culture Collection) planktonic forms treated with different concentrations of Tween 20 [(%)(v/v)] in TSB (Tryptic Soy Broth) medium with regard to untreated cells. Error lines represent the the standard error of the mean (n=6). A red line marks untreated cells.

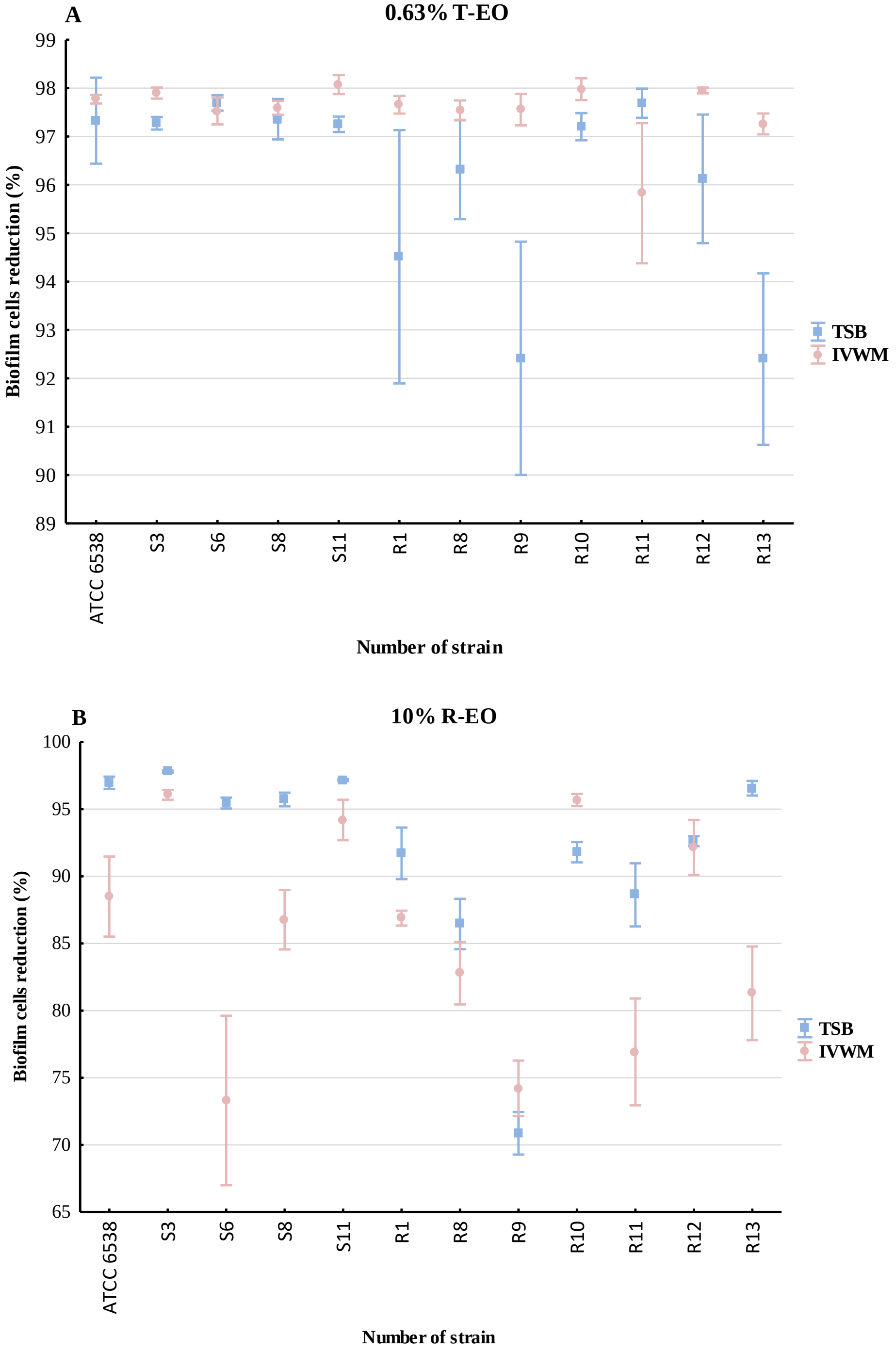

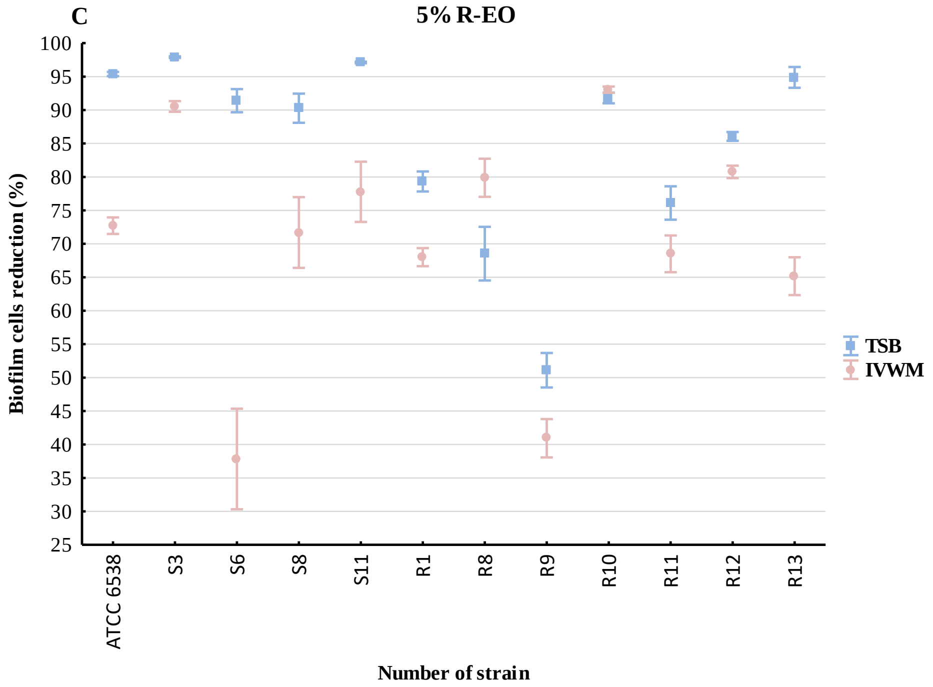
C
5% R-EO
Biofilm cells reduction (%)
100
95
90
85
80
75
70
65
60
55
50
45
40
35
30
25
ATCC 6538
S3
S6
S8
S11
R1
R8
R9
R10
R11
R12
R13
Number of strain
TSB
IVWM

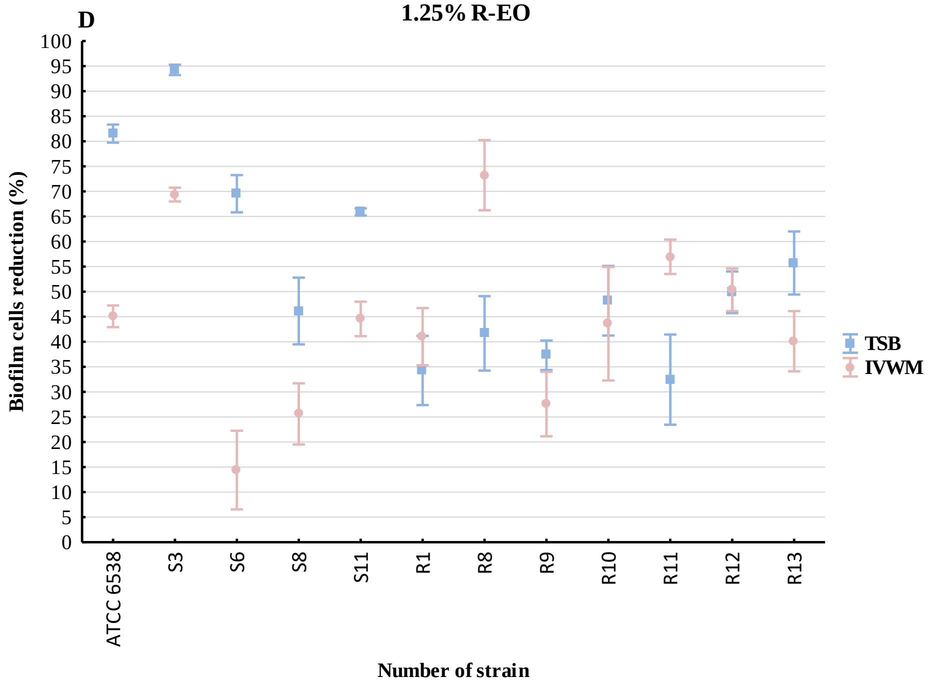
D
1.25% R-EO
Biofilm cells reduction (%)
100
95
90
85
80
75
70
65
60
55
50
45
40
35
30
25
20
15
10
5
0
ATCC 6538
S3
S6
S8
S11
R1
R8
R9
R10
R11
R12
R13
Number of strain
TSB
IVWM

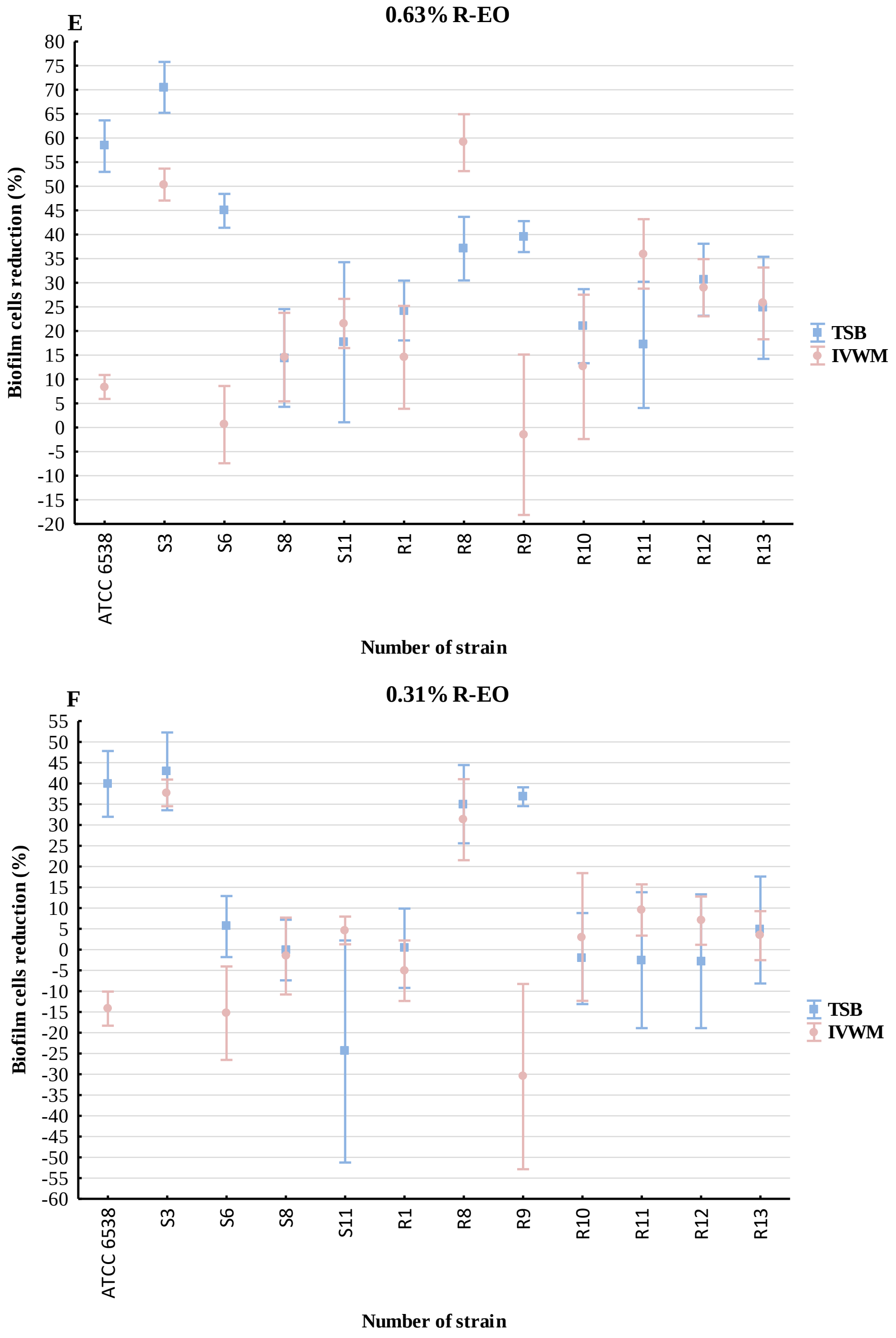


**Supplementary Figure 4**. Average reduction (%) of biofilm cells of reference (ATCC 6538, American Type Culture Collection) or clinical (S3, S6, S8, S11, R1, R8-R13) strains of *S. aureus* after treatment with selected concentrations (%) (v/v) of thyme oil (**A**, T-EO) or rosemary oil (**B-F**, R-EO). Error lines represent the standard error of the mean (n=6). TSB- Tryptic Soy Broth, IVWM- In Vitro Wound Milieu.

### Normal probability plot
### Biofilm biomass TSB

A

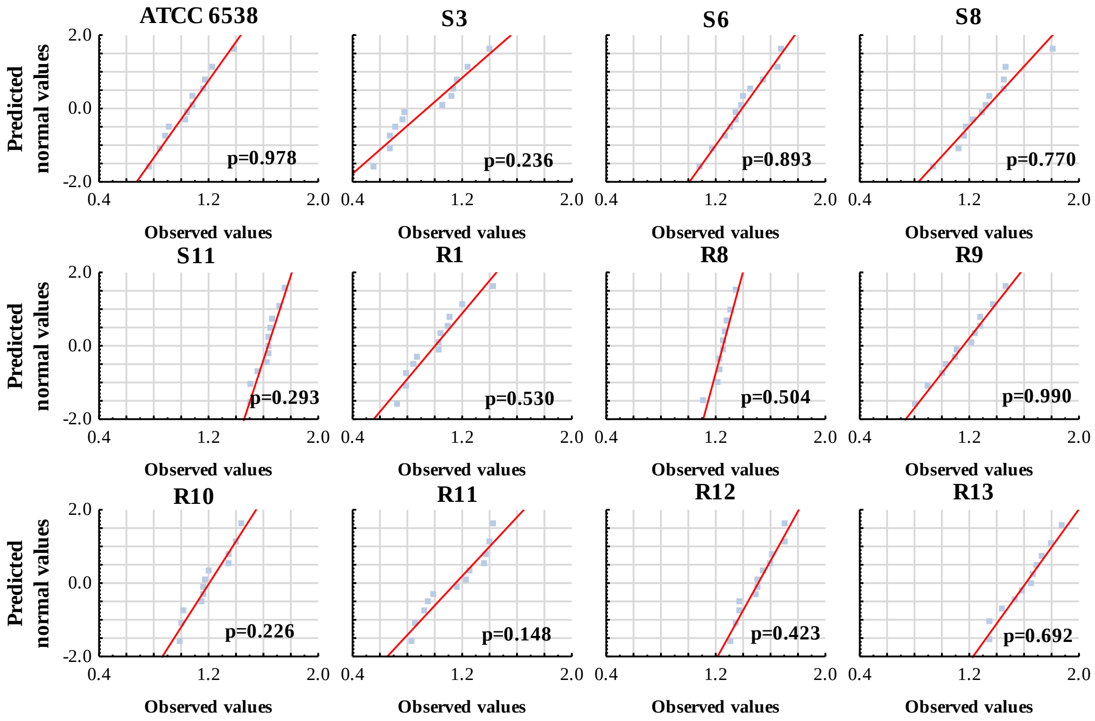


### Normal probability plot
### Biofilm biomass IVWM

B

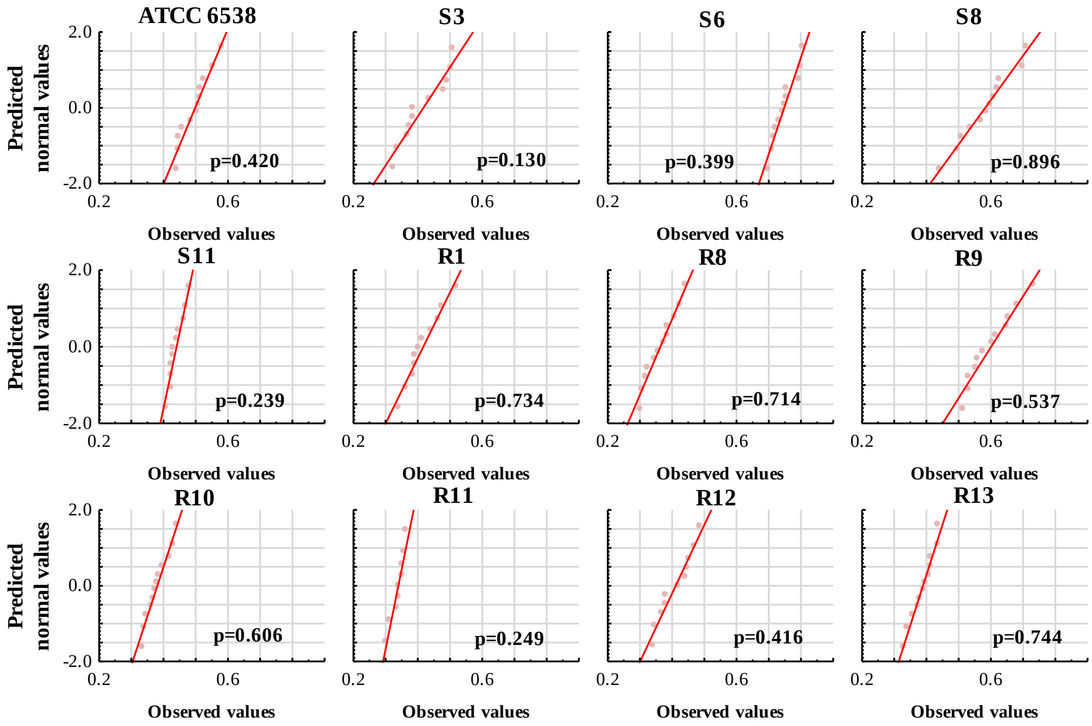

**Supplementary Figure 5**. Normal probability plots for analysis of differences in particular staphylococcal reference (ATCC 6538, American Type Culture Collection) or clinical (S3, S6, S8, S11, R1, R8-R13) strains' ability to form biofilm biomass between Tryptic Soy Broth (**A**, TSB) or In Vitro Wound Milieu (**B**, IVWM) medium. Red curves indicate the matching of the values to the normal distribution. P- probability level calculated with Shapiro-Wilk test. Normal distribution was considered for values of $p>0.05$.

**Supplementary Table 3.** Parameters of the tested statistic of the differences in particular staphylococcal reference (ATCC 6538, American Type Culture Collection) or clinical (S3, S6, S8, S11, R1, R8-R13) strains' ability to form biofilm biomass between Tryptic Soy Broth (TSB) or In Vitro Wound Milieu (IVWM) medium. The normal distribution of the absorbance values was determined in each group (Shapiro-Wilk test, $p<0.05$). Depending on the variance homogeneity (Levene's test, $p<0.05$), t-test (for homogeneous variances, ATCC 6538, S8, S11, R1, R8, R9, R12 strains), or Welch's t-test (for non-homogeneous variances, S3, S6, R10, R11, R13 strains) were performed. T- values of t-test, df- degrees of freedom, p- probability level (values of $p<0.05$ were considered significant).

| **Biofilm biomass** | | | | | |
|---|---|---|---|---|---|
| **Number of strain** | **Average TSB** | **Average IVWM** | **t** | **df** | **p** |
| **ATCC 6538** | 1.1 | 0.4 | 11.6 | 22 | 0.000000 |
| **S3** | 0.9 | 0.2 | 7.9 | 16 | 0.000001 |
| **S6** | 1.4 | 0.9 | 9.0 | 15 | 0.000000 |
| **S8** | 1.3 | 0.6 | 9.6 | 22 | 0.000000 |
| **S11** | 1.6 | 0.3 | 50.0 | 20 | 0.000000 |
| **R1** | 1.0 | 0.2 | 11.2 | 21 | 0.000000 |
| **R8** | 1.3 | 0.1 | 32.0 | 20 | 0.000000 |
| **R9** | 1.2 | 0.6 | 8.1 | 22 | 0.000000 |
| **R10** | 1.2 | 0.2 | 21.3 | 15 | 0.000000 |
| **R11** | 1.2 | 0.1 | 16.2 | 12 | 0.000000 |
| **R12** | 1.5 | 0.2 | 25.7 | 21 | 0.000000 |
| **R13** | 1.6 | 0.2 | 25.3 | 13 | 0.000000 |

### Normality distribution

### Biofilm metabolic activity TSB

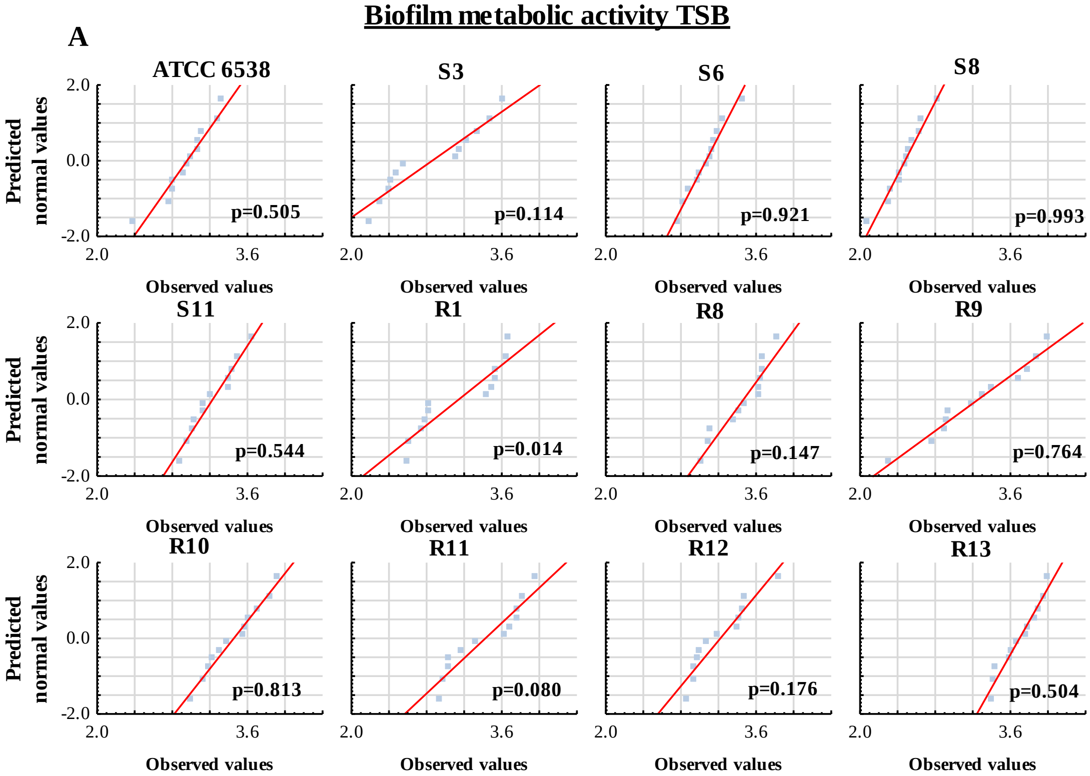


### Normality distribution

### Biofilm metabolic activity IVWM

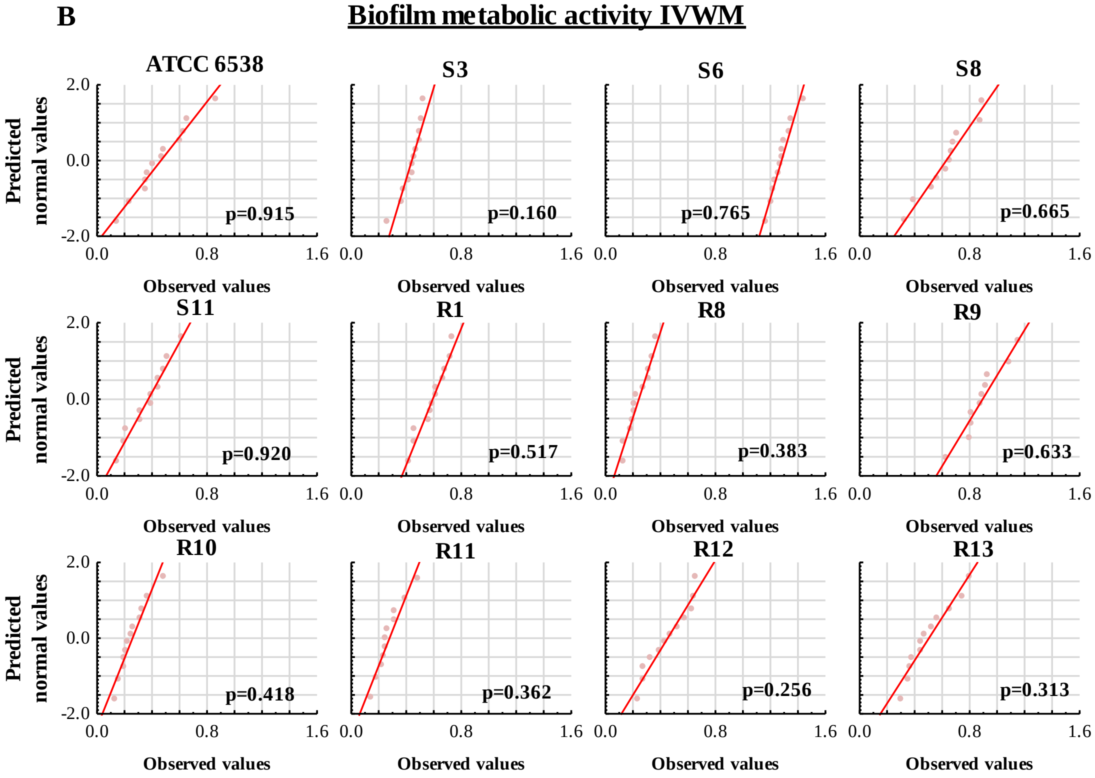

**Supplementary Figure 6**. Normal probability plots for analysis of differences in particular staphylococcal reference (ATCC 6538, American Type Culture Collection) or clinical (S3, S6, S8, S11, R1, R8-R13) strains' metabolic activity of biofilms between Tryptic Soy Broth (**A**, TSB) or In Vitro Wound Milieu (**B**, IVWM) medium. Red curves indicate the fitting of the values to the normal distribution. P- probability level calculated with Shapiro-Wilk test. Normal distribution was considered for values of $p>0.05$.

**Supplementary Table 4.** Parameters of the tested statistic of the differences in particular staphylococcal reference (ATCC 6538, American Type Culture Collection) or clinical (S3, S6, S8, S11, R1, R8-R13) strains' metabolic activity of biofilms between Tryptic Soy Broth (TSB) or In Vitro Wound Milieu (IVWM) medium. Normal distribution of the values was determined in each group except the R1 strain (Shapiro-Wilk test, $p<0.05$). Depending on the variance homogeneity (Levene's test, $p<0.05$), t-test (for homogeneous variances, ATCC 6538, S8, R13 strains) or Welch's t-test (for non-homogeneous variances, S3, S6, S11, R8, R9, R10, R11, R12 strains) were performed in groups with the normal distribution of values. For the R1 strain, Mann-Whitney U test was performed. T-values of t-test, df- degrees of freedom, p- probability level (values of $p<0.05$ were considered significant), U and Z- values of U-test.

| **Biofilm metabolic activity** | | | | | |
|---|---|---|---|---|---|
| **Number of strain** | **Average TSB** | **Average IVWM** | **t** | **df** | **p** |
| **ATCC 6538** | 3.0 | 0.5 | 26.7 | 22 | 0.000000 |
| **S3** | 2.9 | 0.4 | 16.2 | 11 | 0.000000 |
| **S6** | 3.1 | 1.3 | 30.2 | 14 | 0.000000 |
| **S8** | 2.5 | 0.6 | 24.3 | 21 | 0.000000 |
| **S11** | 3.2 | 0.4 | 35.4 | 18 | 0.000000 |
| **R8** | 3.5 | 0.2 | 40.4 | 13 | 0.000000 |
| **R9** | 3.3 | 0.9 | 15.2 | 13 | 0.000000 |
| **R10** | 3.5 | 0.3 | 35.9 | 13 | 0.000000 |
| **R11** | 3.4 | 0.3 | 27.7 | 13 | 0.000000 |
| **R12** | 3.2 | 0.5 | 28.7 | 16 | 0.000000 |
| **R13** | 3.7 | 0.5 | 42.0 | 22 | 0.000000 |
| | **Rank sum TSB** | **Rank sum IVWM** | **U** | **Z** | **p** |
| **R1** | 222 | 78 | 0 | 4 | 0.000037 |

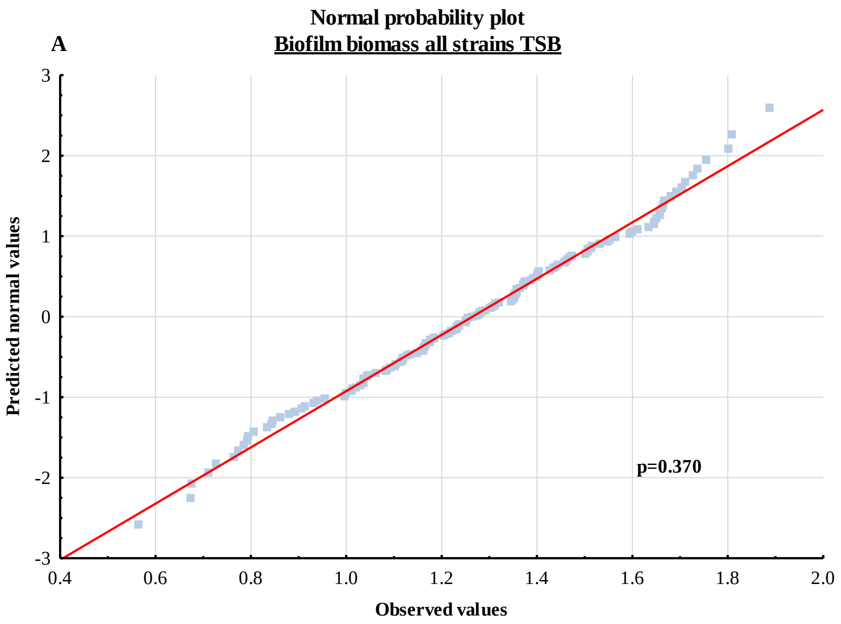
Normal probability plot
Biofilm biomass all strains TSB
A
Predicted normal values
Observed values
p=0.370
3
2
1
0
-1
-2
-3
0.4
0.6
0.8
1.0
1.2
1.4
1.6
1.8
2.0

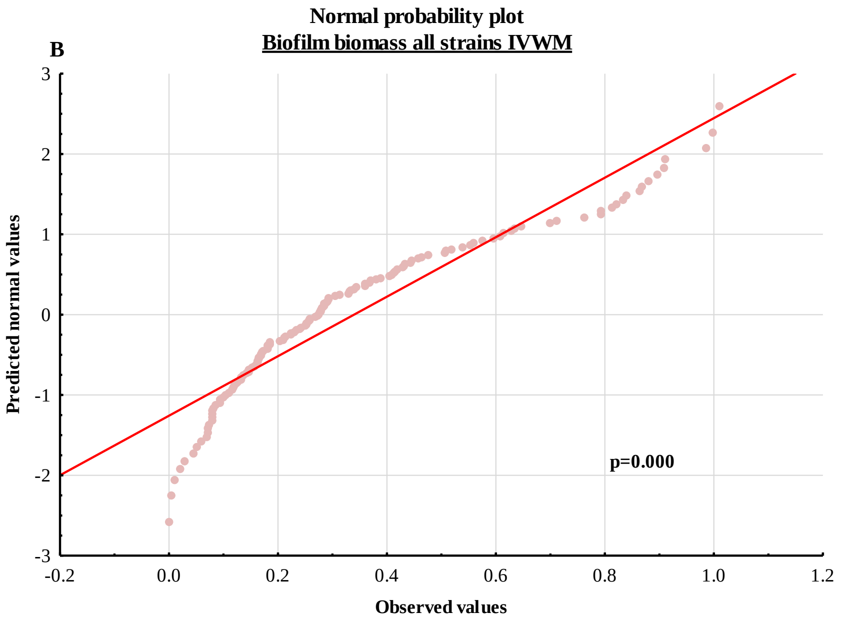
Normal probability plot
Biofilm biomass all strains IVWM
B
Predicted normal values
Observed values
p=0.000
3
2
1
0
-1
-2
-3
-0.2
0.0
0.2
0.4
0.6
0.8
1.0
1.2

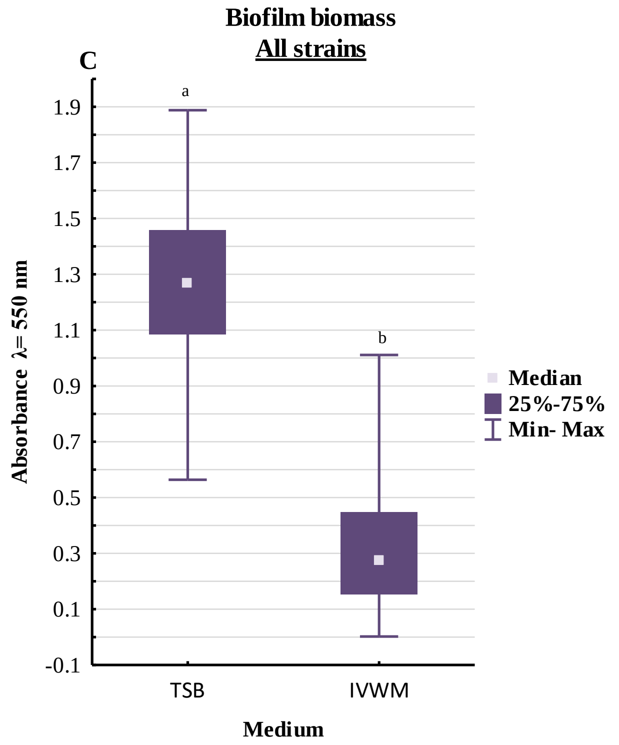
Biofilm biomass
All strains
C
a
b
1.9
1.7
1.5
1.3
1.1
0.9
0.7
0.5
0.3
0.1
-0.1
Absorbance λ= 550 nm
Median
25%-75%
Min- Max
TSB
IVWM
Medium

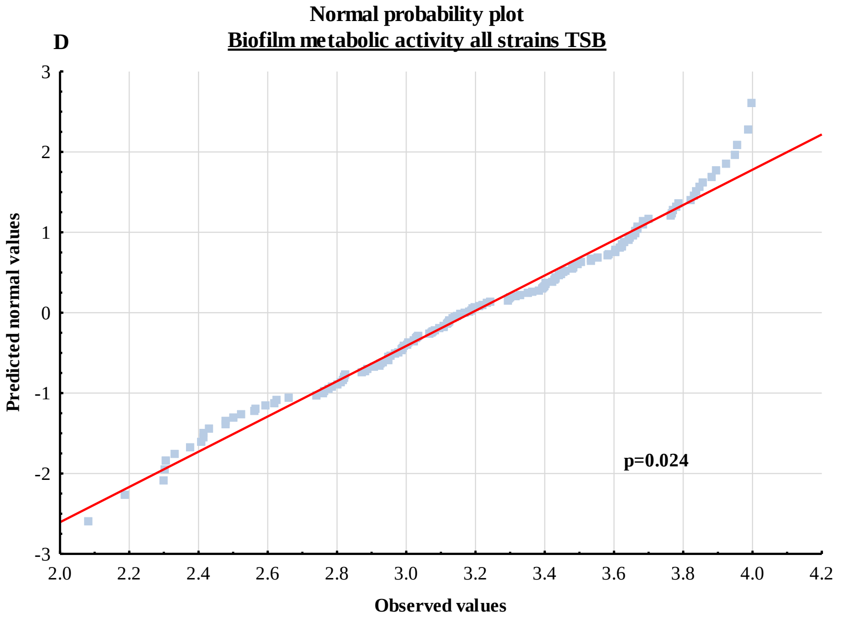
Normal probability plot
Biofilm metabolic activity all strains TSB
D
3
2
1
0
-1
-2
-3
Predicted normal values
p=0.024
2.0
2.2
2.4
2.6
2.8
3.0
3.2
3.4
3.6
3.8
4.0
4.2
Observed values

E

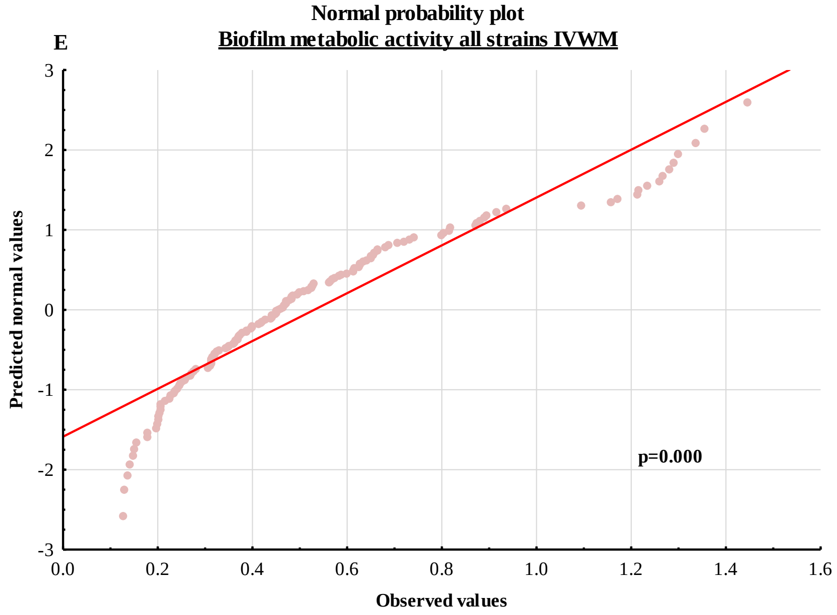


F

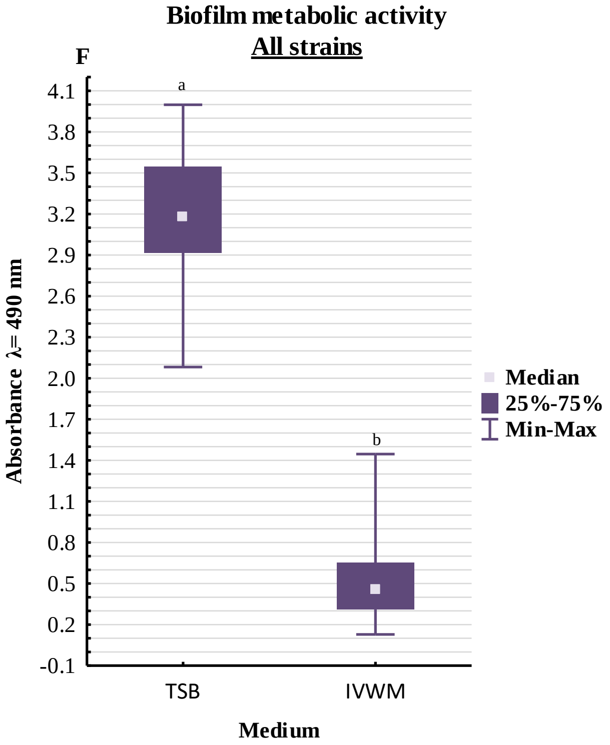


**Supplementary Figure 7**. Detailed analysis of differences in all (as a group) staphylococcal strains' ability to form biofilm biomass or their metabolic activity between Tryptic Soy Broth (TSB) or In Vitro Wound Milieu (IVWM) medium. Normal probability plots for biofilm biomass in TSB (**A**) or IVWM (**B**) medium. Box and whisker plot of the absorbance values of biofilm biomass (**C**). Normal probability plots for biofilm metabolic activity in TSB (**D**) or IVWM (**E**) medium. Box and whisker plot of the absorbance values of biofilm metabolic activity (**F**). 25%-75%- interquartile range, Min-Max- a range between minimal and maximum values. Red curves indicate the fitting of the values to the normal distribution. The statistically significant differences are marked with pairs of letters a/b. P-probability level calculated with Shapiro-Wilk test. Normal distribution was considered for values of $p>0.05$.

**Supplementary Table 5.** Parameters of the tested statistics of the differences in all (as a group) staphylococcal strains' ability to form biofilm biomass or their metabolic activity between Tryptic Soy Broth (TSB) or In Vitro Wound Milieu (IVWM) medium. Non-parametric Mann-Whitney U test was performed because the distributions were non-normal strain (Shapiro-Wilk test, $p<0.05$). P- probability level (values of $p<0.05$ were considered significant), U and Z- values of U-test.

| Analysis of all strains as a group | | | | | |
|---|---|---|---|---|---|
| **Type of compared biofilm characteristic** | **Rank sum TSB** | **Rank sum IVWM** | **U** | **Z** | **p** |
| **Biofilm biomass** | 28802 | 9702 | 249 | 14 | 0.000000 |
| **Biofilm metabolic activity** | 30600 | 9870 | 0 | 15 | 0.000000 |

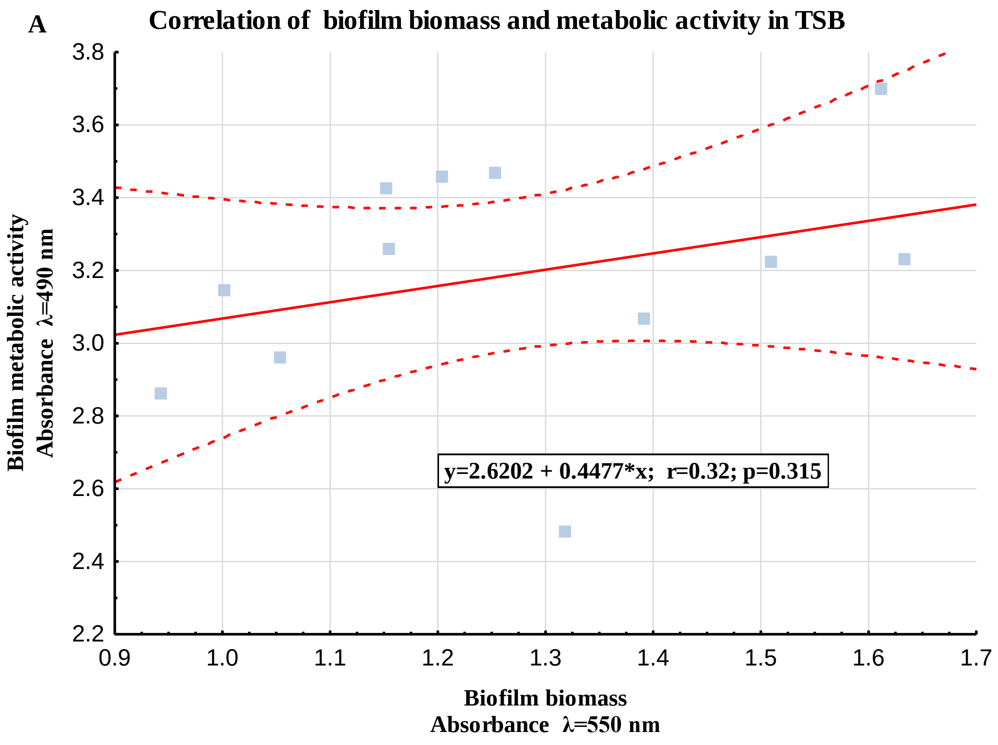

B

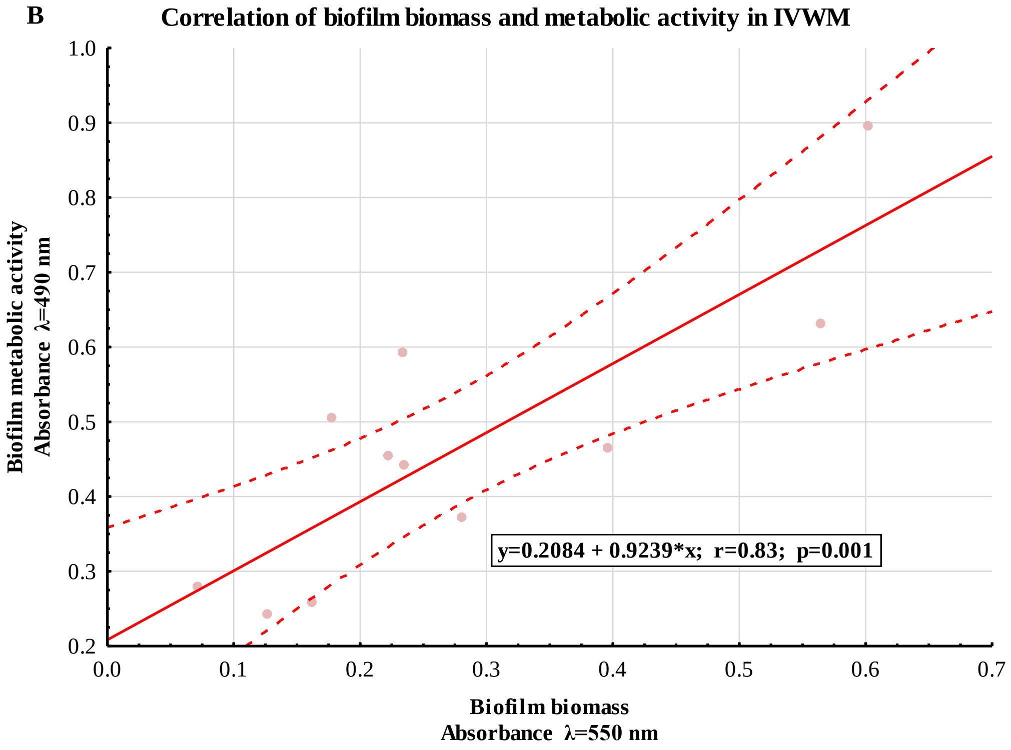


**Supplementary Figure 8.** Scatter plots of correlations of average staphylococcal strains' ability to form biofilm biomass and average metabolic activity for strains cultured in Tryptic Soy Broth (TSB, **A**) or In Vitro Wound Milieu (IVWM, **B**) medium. R- correlation coefficient, p- probability level (values of $p<0.05$ were considered significant). Red lines indicate 95% confidence intervals.

**Supplementary Table 6.** Results of the correlation analysis of average staphylococcal strains' ability to form biofilm biomass and average metabolic activity for strains cultured in Tryptic Soy Broth (TSB) or In Vitro Wound Milieu (IVWM) medium. Avr- average, SD- standard deviation, r- correlation coefficient, t- values of statistics, p- probability level (values of $p<0.05$ were considered significant).

| | Correlation of biofilm biomass and metabolic activity | | | | | | |
|---|---|---|---|---|---|---|---|
| | Biofilm biomass | | Metabolic activity | | | | |
| **Medium** | **Avr** | **SD** | **Avr** | **SD** | **r** | **t** | **p** |
| **TSB** | 1.3 | 0.2 | 3.2 | 0.3 | 0.32 | 1.1 | 0.314698 |
| **IVWM** | 0.3 | 0.2 | 0.5 | 0.2 | 0.83 | 4.5 | 0.001463 |

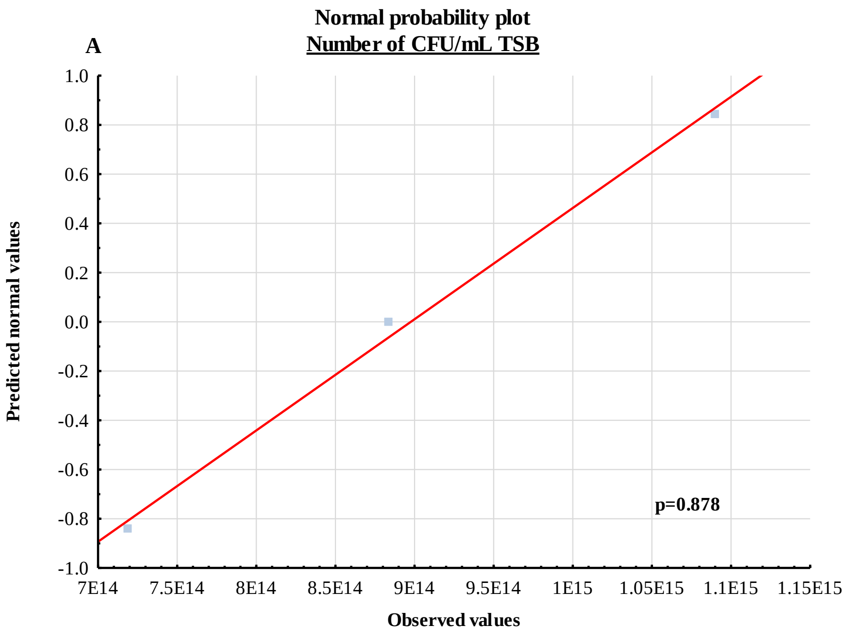
A
Normal probability plot
Number of CFU/mL TSB
1.0
0.8
0.6
0.4
0.2
0.0
-0.2
-0.4
-0.6
-0.8
-1.0
Predicted normal values
7E14
7.5E14
8E14
8.5E14
9E14
9.5E14
1E15
1.05E15
1.1E15
1.15E15
Observed values
p=0.878

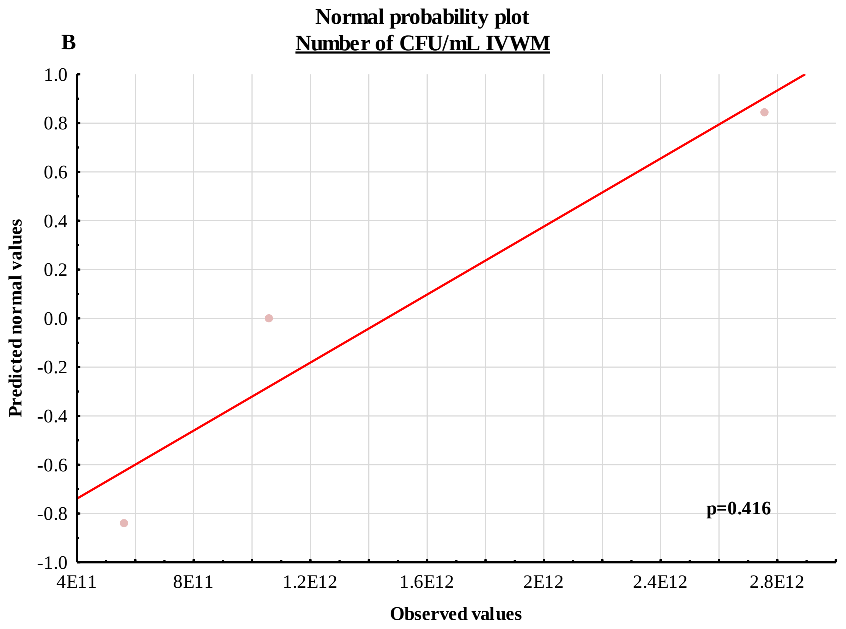
B
Normal probability plot
Number of CFU/mL IVWM
1.0
0.8
0.6
0.4
0.2
0.0
-0.2
-0.4
-0.6
-0.8
-1.0
Predicted normal values
4E11
8E11
1.2E12
1.6E12
2E12
2.4E12
2.8E12
Observed values
p=0.416

C

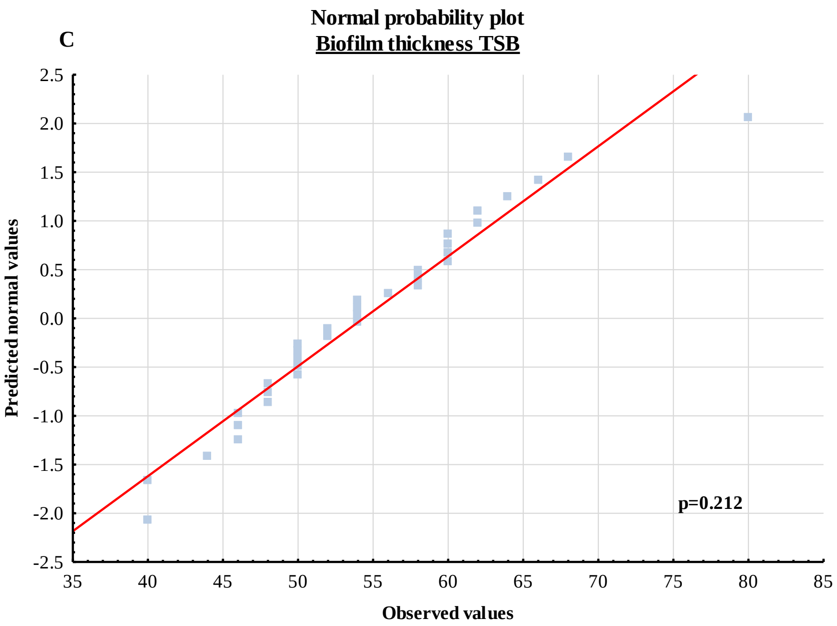
Normal probability plot
Biofilm thickness TSB
Predicted normal values
Observed values
p=0.212

D

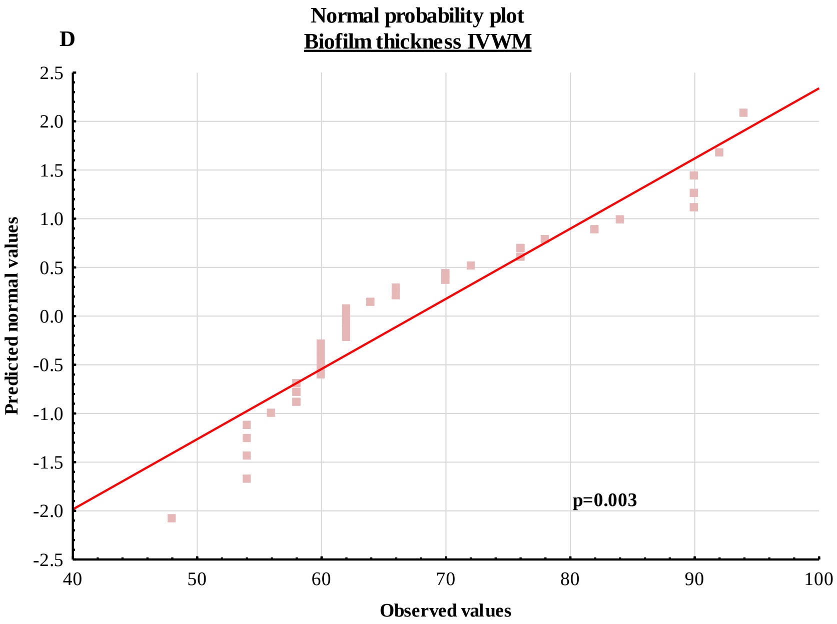
Normal probability plot
Biofilm thickness IVWM
Predicted normal values
Observed values
p=0.003

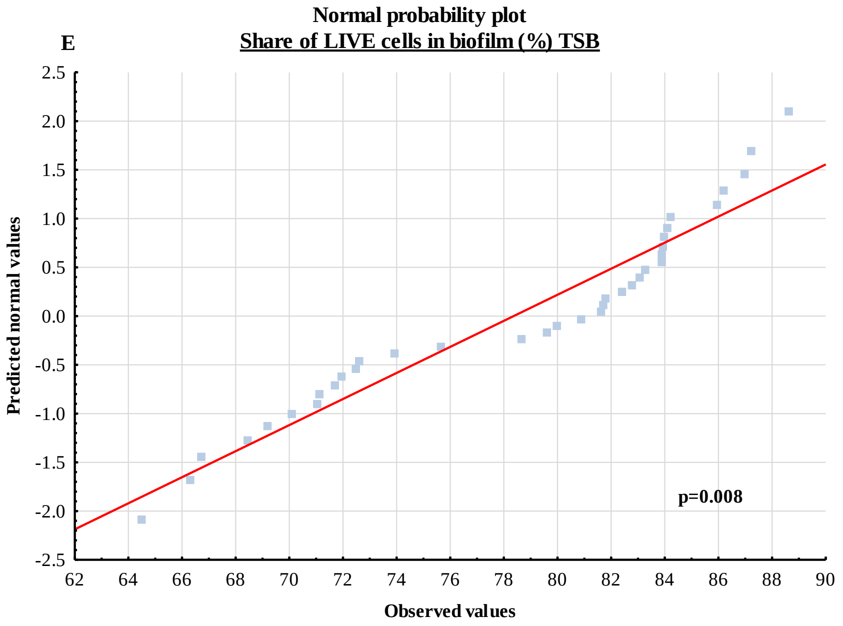
E
Normal probability plot
Share of LIVE cells in biofilm (%) TSB
Predicted normal values
Observed values
p=0.008
2.5
2.0
1.5
1.0
0.5
0.0
-0.5
-1.0
-1.5
-2.0
-2.5
62 64 66 68 70 72 74 76 78 80 82 84 86 88 90

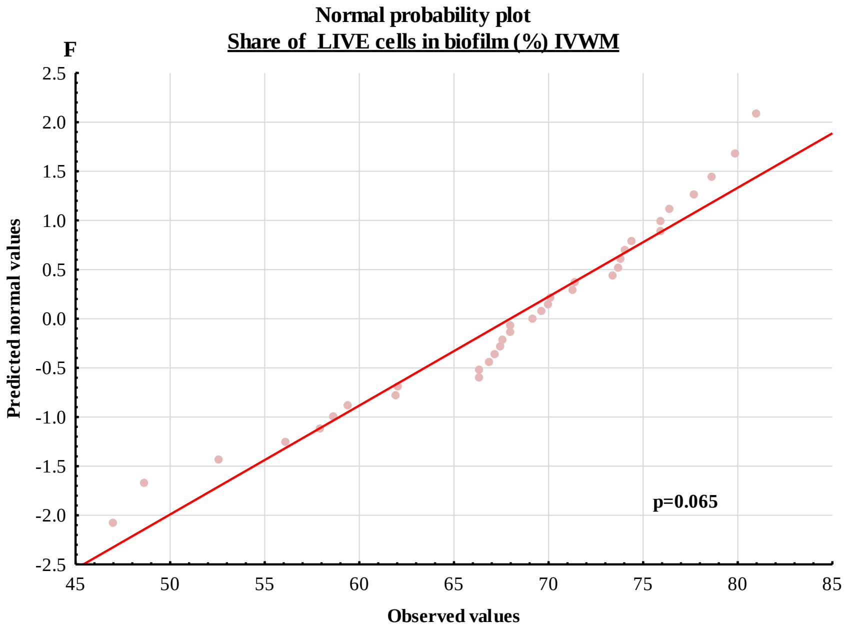
F
Normal probability plot
Share of LIVE cells in biofilm (%) IVWM
Predicted normal values
Observed values
p=0.065
2.5
2.0
1.5
1.0
0.5
0.0
-0.5
-1.0
-1.5
-2.0
-2.5
45 50 55 60 65 70 75 80 85

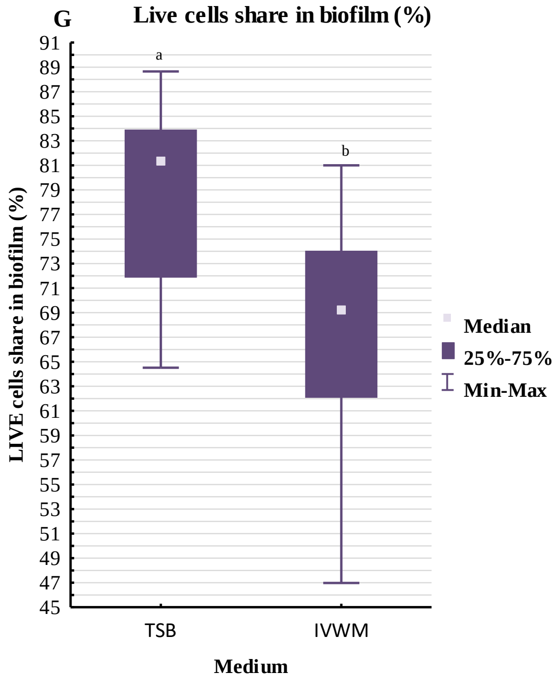


**Supplementary Figure 9**. Detailed analysis of the differences in the number of CFU/mL (Colony-Forming Units), biofilm thickness, and percentage share of LIVE cells between staphylococcal biofilms cultured in Tryptic Soy Broth (TSB) or In Vitro Wound Milieu (IVWM) medium. Normal probability plots for the number of CFU/mL in TSB (**A**) or IVWM (**B**) medium. Normal probability plots for biofilm thickness in TSB (**C**) or IVWM (**D**) medium. Normal probability plots for a share of LIVE cells in biofilm in TSB (**E**) or IVWM (**F**) medium. Box and whisker plot of share of LIVE cells in biofilm (**G**). 25%-75%- interquartile range, Min-Max- a range between minimal and maximum values. Red curves indicate the fitting of the values to the normal distribution. The statistically significant difference is marked with a pair of letters a/b. P- probability level calculated with Shapiro-Wilk test. Normal distribution was considered for values of $p>0.05$.

**Supplementary Table 7.** Parameters of the tested statistics of the differences in the number of CFU/mL (Colony-Forming Units), biofilm thickness, and percentage share of LIVE cells between staphylococcal strains cultured in Tryptic Soy Broth (TSB) or In Vitro Wound Milieu (IVWM) medium. Normal distribution of the values and homogeneity of variances were determined in the number of CFU/mL (Shapiro-Wilk test, Levene's test, respectively $p<0.05$), therefore t-test was performed. Mann-Whitney U test was performed for biofilm thickness and share of LIVE cells because the distributions were non-normal. T- values of t-test, df- degrees of freedom, p- probability level (values of $p<0.05$ were considered significant), U and Z- values of U-test.

| **Analysis of other biofilm characteristics** | | | | | |
|---|---|---|---|---|---|
| **Type of compared characteristic** | **Average TSB** | **Average IVWM** | **t** | **df** | **p** |
| **Number of CFU/mL** | 9.0E+14 | 1.5E+12 | 8.4 | 4 | 0.001124 |
| | **Rank sum TSB** | **Rank sum IVWM** | **U** | **Z** | **p** |
| **Biofilm thickness** | 813 | 1602 | 218 | -5 | 0.000006 |
| **Share of LIVE cells** | 1702 | 854 | 224 | 5 | 0.000003 |

**Supplementary Table 8.** Results of the multivariate analysis of variance for biofilm cells reduction after T-EO treatment. F- values of the test, p- probability level (values of p<0.05 were considered significant).

| Effect | f | p |
|---|---|---|
| **Medium** | 13.7 | 0.000239 |
| **Strain** | 1.9 | 0.036016 |
| **T-EO concentration (%)** | 701.9 | 0.000000 |
| **Medium and Strain** | 4.0 | 0.000013 |
| **Medium and T-EO concentration (%)** | 10.9 | 0.000000 |
| **Strain and T-EO concentration (%)** | 1.8 | 0.002131 |
| **Medium and Strain and T-EO concentration (%)** | 3.7 | 0.000000 |

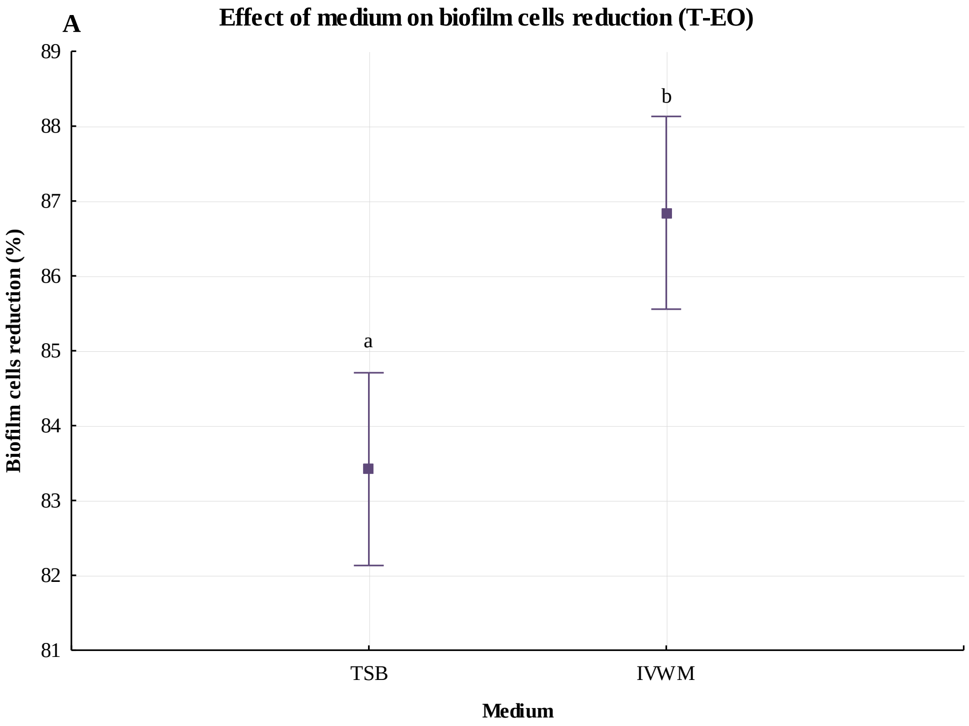

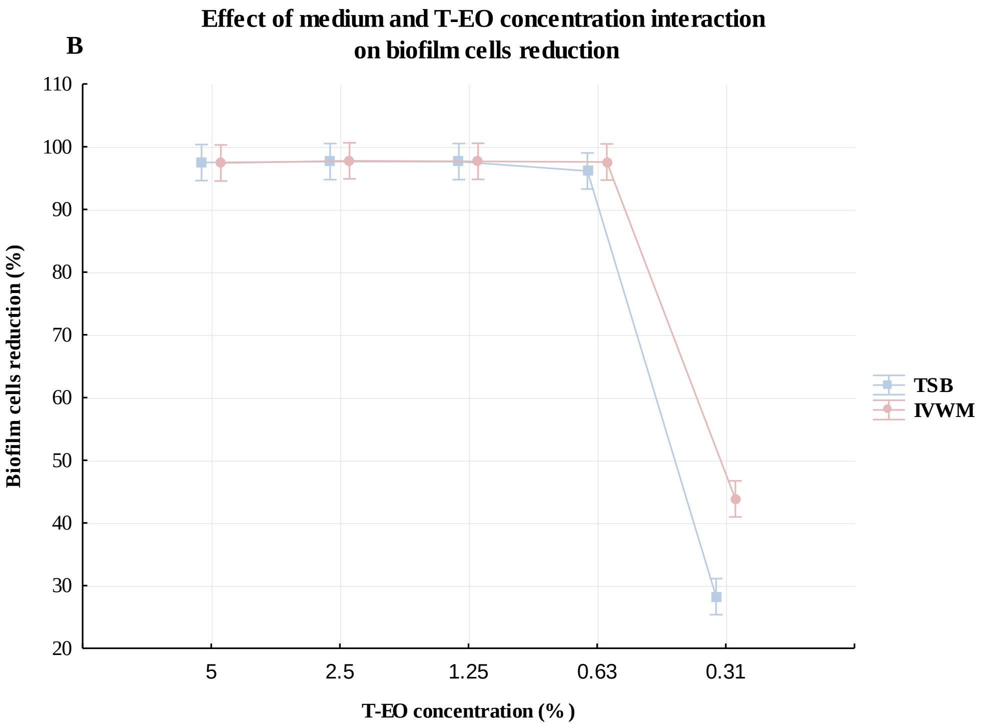


**Supplementary Figure 10.** Effect of medium (**A**) or medium and EO concentration (%) (v/v) interaction (**B**) on biofilm cells reduction after treatment with T-EO (thyme oil). Results of the multivariate analysis of variance. The statistically significant difference is marked with pairs of letters a/b. Tags denote means, and vertical bars denote 0.95 confidence intervals. Tags are dislocated on the X-axis values to gain higher visibility. TSB- Tryptic Soy Broth, IVWM- In Vitro Wound Milieu.

**Supplementary Table 9.** Results of the multivariate analysis of variance for biofilm cells reduction after R-EO treatment. F- values of the test, p- probability level (values of $p<0.05$ were considered significant).

| Effect | f | p |
|---|---|---|
| **Medium** | 83.9 | 0.000000 |
| **Strain** | 30.0 | 0.000000 |
| **R-EO concentration (%)** | 488.2 | 0.000000 |
| **Medium and Strain** | 17.1 | 0.000000 |
| **Medium and R-EO concentration (%)** | 1.1 | 0.379590 |
| **Strain and R-EO concentration (%)** | 3.1 | 0.000000 |
| **Medium and Strain and R-EO concentration (%)** | 2.6 | 0.000000 |

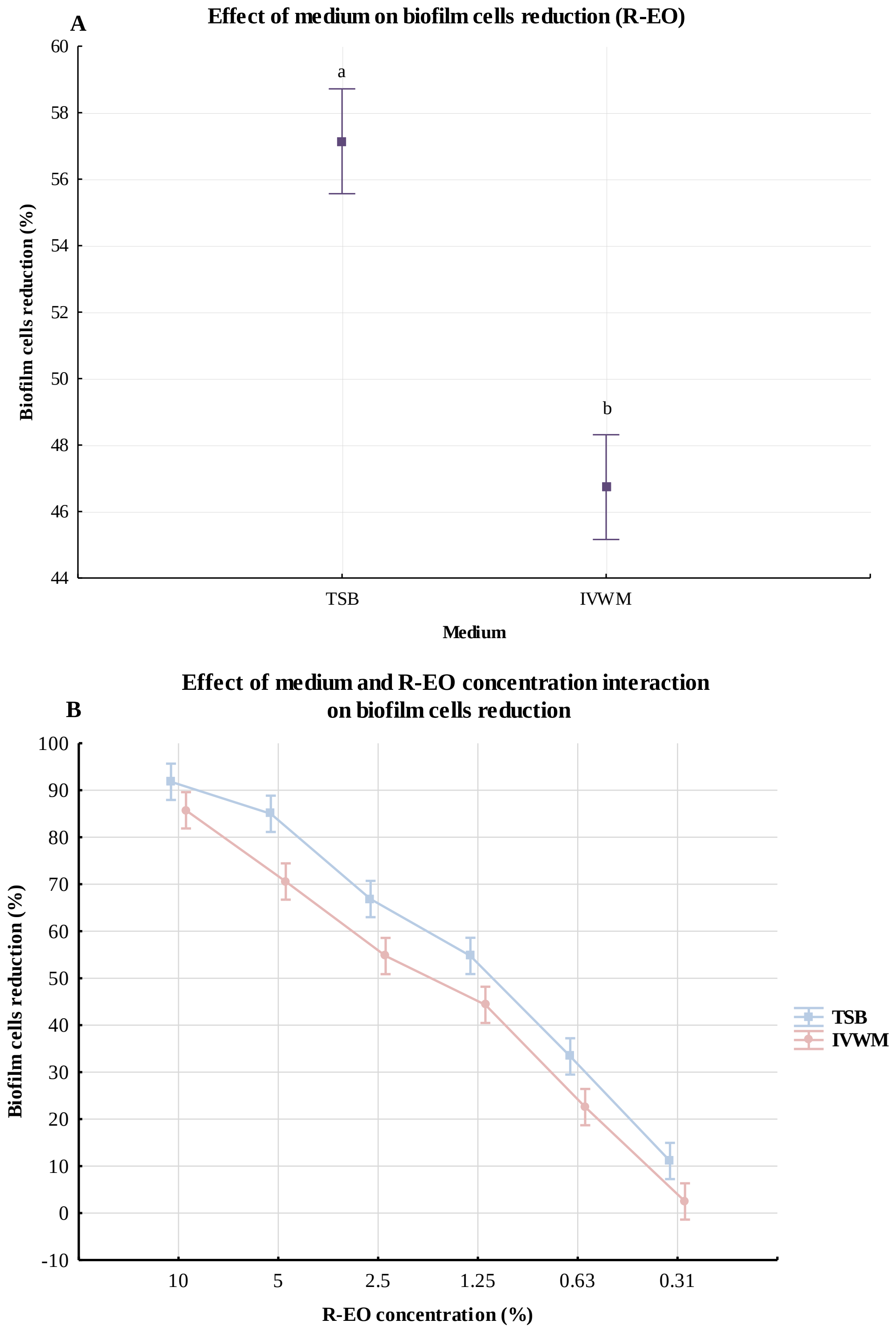


**Supplementary Figure 11.** Effect of medium (**A**) or medium and EO concentration (%) (v/v) interaction (**B**) on biofilm cells reduction after treatment with R-EO (rosemary oil). Results of the multivariate analysis of variance. Tags denote means, and vertical bars denote 0.95 confidence intervals. The statistically significant difference is marked with pairs of letters a/b. Tags are dislocated on the X-axis values to gain higher visibility. TSB- Tryptic Soy Broth, IVWM- In Vitro Wound Milieu.
